# Spatial growth within food-like matrices drives physiological diversification and stress adaptation in *Salmonella*

**DOI:** 10.64898/2026.11.16.766896

**Authors:** Elodie Hoch, Yasmine Dergham, Anaïs Pierron, Julien Deschamps, Florence Dubois-Brissonnet, Jérôme Lecardonnel, Hans Steenackers, Dalal Werner, Bernard Hezard, Claudia Bevilacqua, Lysiane Omhover-Fougy, Romain Briandet

## Abstract

Foodborne bacteria often grow in structured environments, but the effects of spatial organisation on their physiology and subsequent stress tolerance remain poorly understood. In this study, we investigated how spatial growth in a food-like hydrogel matrix affected the physiology of *Salmonella enterica* serovar Typhimurium and its ability to cope with environmental stresses. Planktonic and spatialised populations were grown for four days in media without NaCl supplementation or supplemented with 6% NaCl. Transcriptomic, extracellular metabolomic, and lipidomic analyses were combined with targeted spatial gene-expression analyses. The multi-omics analyses showed that spatial organisation had a strong effect on *Salmonella* physiology and induce more physiological variation than the presence of osmotic stress, under the tested conditions. Spatial growth was associated with significant changes in gene expression, carbon metabolism, and membrane fatty-acid composition. Several functions related to stress response and biofilm-associated behaviour were also affected, with corresponding genes showing significant differential expression. When bacteria were grown in the presence of NaCl, spatialised populations showed less transcriptional and metabolic reorganisation than planktonic populations. In addition, gene-expression analyses showed that microcolonies were physiologically heterogeneous, with differences among cell populations within the same microcolony. Spatial growth also associated with the upregulation of functions related to the general stress response, oxidative stress protection, membrane adaptation and acid stress. These physiological changes provide a plausible mechanistic framework for the enhanced gastrointestinal stress tolerance previously observed in spatially grown populations, particularly under acid challenge. Overall, our results show that spatial organisation is an important environmental factor that reprograms *Salmonella* physiology and can influence its ability to survive stresses encountered after food ingestion.

## Introduction

Bacteria rarely grow in homogeneous environments. In natural ecosystems, host-associated niches and food matrices, bacterial cells frequently experience spatial constraints that influence nutrient availability, oxygen diffusion, metabolite accumulation, and cell–cell interactions. These local conditions can generate physiologically distinct populations within genetically identical communities and strongly influence bacterial behaviour, including growth, stress response, and survival (**Flemming et al. 2016; Jo, Price-Whelan, Dietrich 2022; Stewart, Franklin 2008**). Therefore, the spatial organisation of bacterial growth is increasingly recognised as an important parameter that controls microbial physiology, alongside the chemical composition of the environment (**Nadell, Drescher, Foster 2016; Nath et al. 2022**). Some studies investigated the role of matrices, and microorganisms confinement in their responses to environmental stresses (**Saint Martin, Jubelin, et al. 2022**), but, most studies investigating the physiology and stress responses of foodborne pathogens free cells grown in homogeneous planktonic media.

Most of our knowledge about spatially organised bacterial populations comes from biofilm studies. In these communities, cells attach to surfaces, produce extracellular matrix components, and progressively generate physicochemical gradients that create heterogeneous metabolic and physiological states (**Dergham et al. 2023; Jo, Price-Whelan, Dietrich 2022; Stewart, Franklin 2008**). Biofilms are therefore not simple aggregates of cells but structured microbial communities in which spatial organisation contributes to adaptation, stress tolerance and survival (**Avcı, Akçelik, Akçelik 2026; Ban-Cucerzan et al. 2025; Hall-Stoodley, Costerton, Stoodley 2004; Fàbrega, Vila 2013**).

Recent studies have further shown that spatially organised bacterial communities are composed of distinct transcriptional states rather than homogeneous populations. In *Bacillus subtilis*, spatially resolved transcriptomic analyses revealed the coexistence of metabolically distinct subpopulations within biofilms, demonstrating that local organisation can generate functional heterogeneity at the microscale (**Dergham et al. 2023**). However, spatial organisation does not only result from genetic modifications related to this lifestyle. Recent studies have highlighted the role of the physical environment itself as a determinant of bacterial physiology. Bacteria can sense and respond to mechanical and spatial constraints through changes in growth, motility and stress responses (**Dufrêne, Persat 2020**). For example, the growth of *Pseudomonas aeruginosa* under mechanical confinement in hydrogels has recently been shown to increase antibiotic tolerance through physiological adaptations involving ion transport and membrane remodelling (**Monnappa et al. 2026**). Other studies, concerning structured populations of *Salmonella* originates from studies of surface-associated biofilms, particularly the red, dry and rough (rdar) phenotype (**Römling 2005)**. This multicellular state is characterised by the production of extracellular matrix components, including curli and cellulose, and is controlled by regulatory networks involving *csgD* and c-di-GMP signalling (**Simm et al. 2014; Solano et al. 2002; Zakikhany et al. 2010**). These studies have demonstrated that *Salmonella* can transition between distinct physiological states during biofilm development, with consequences for stress tolerance and persistence in different environments. Finally, these observations demonstrate that the physical context in which bacteria grow can influence their subsequent response to stress. However, surface-associated biofilms represent only one form of bacterial spatial organisation (**Jeanson et al. 2015; Serra, Hengge 2014**).

Food matrices represent particularly relevant examples of structured bacterial habitats. Unlike homogeneous laboratory cultures, foods are heterogeneous three-dimensional environments in which bacteria can become locally immobilised and develop as spatially organised microcolonies. Such an organisation may influence bacterial physiology by modifying diffusion processes, local nutrient availability and interactions between neighbouring cells (**Nath et al. 2022**). Hydrogel-based models that reproduce the physical properties of structured foods provide a controlled approach to investigating these processes and this spatial growth modifies bacterial behaviour (Saint Martin, Darsonval, et al. 2022). In *Escherichia coli* O157:H7, growth within structured matrices generated spatial heterogeneity in the expression of the acid-resistance gene *gadB*, with higher expression detected in specific regions of microcolonies (**Saint Martin et al. 2023**). During food contamination, *Salmonella* can also grow within three-dimensional structured matrices where bacterial localisation is imposed by the surrounding environment rather than initiated by surface attachment. Recently, we showed that spatial growth in food-like matrices altered responses to food-related stresses and increased subsequent gastrointestinal tolerance in several major foodborne pathogens, including *Salmonella* Typhimurium (**Hoch et al. 2026**). These observations suggest that the spatial organisation experienced during growth can influence bacterial physiology beyond the immediate environmental conditions. However, the mechanisms responsible for this effect remain unknown.

Here, we investigated how growth as spatially organised microcolonies within a food-like matrix modifies the physiology of *S. enterica* serovar Typhimurium and its response to osmotic stress. Among food-associated stresses, osmotic stress is one of the most frequently encountered by foodborne pathogens. Salts remains widely used in the food industry as a preservation strategy in products such as cured meats, cheeses and fermented food. According to our previous study (**Hoch et al. 2026**), a concentration of 6% NaCl show a strong differential impact on *Salmonella* growth between planktonic and spatialised lifestyles. To correlate this phenotypic characteristic to molecular signatures, we compared planktonic populations and matrix-embedded clonal microcolonies grown in the presence or absence of NaCl using integrated transcriptomic, metabolomic, and lipidomic approaches. We further investigated whether the physiological state associated with spatial growth was homogeneous within individual microcolonies using spatial gene-expression analyses.

## Results

### 1. Spatial structuring is the major driver of multi-omics variability in *Salmonella*

To obtain an integrated overview of the physiological impact of spatial structuring and osmotic stress, transcriptomic, metabolomic and lipidomic datasets were analysed simultaneously using DIABLO framework of the MixOmics R package (**Figure 1**). The three omics datasets revealed a clear separation of the four experimental conditions: planktonic populations without NaCl (PT), planktonic populations supplemented with NaCl (PNaCl), spatialised populations without NaCl supplementation (ST), and spatialised populations supplemented with NaCl (SNaCl), indicating that both lifestyle and osmotic stress affect *Salmonella* physiology (**Figure 1A**). Within each omics block, biological replicates clustered tightly, demonstrating high reproducibility of the datasets. To obtain a global view of the physiological responses upregulated by the lifestyle (planktonic versus spatialised in hydrogel) and osmotic stress, transcriptomic, metabolomic and lipidomic datasets were integrated into a single PCA (see **Supplementary Figure S1).** This representation included a total of 4,645 variables measured across the four experimental conditions (PT, PNaCl, ST and SNaCl). The PCA revealed a clear separation of samples by lifestyle along PC1, which accounted for 39.8% of the total variance. By contrast, the effect of osmotic stress was captured mainly by PC2, which explained 25.9% of the variance. Replicates clustered tightly within each condition, confirming the high reproducibility of the multi-omics datasets.

**Figure 1:**
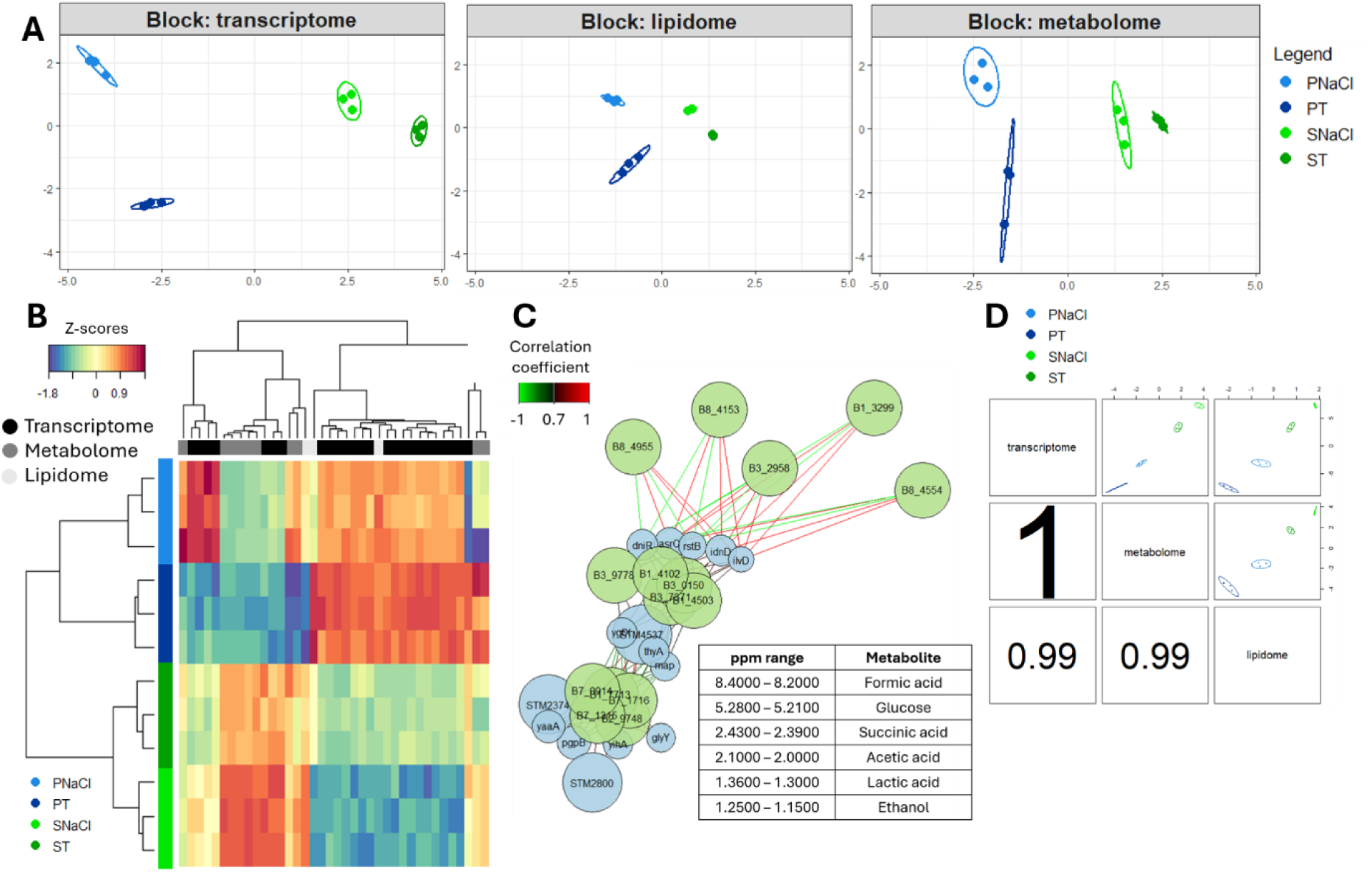
DIABLO (multi-block sPLSDA) integrating multi-omics data. **A**: Plot showing the individual samples from the three omics datasets used in the DIABLO analysis, with different colours representing distinct sample groups (light blue: PNaCl; dark blue: PT; light green: SNaCl; and dark green: ST). **B**: Clustered image map (CIM) (Euclidean distance, complete linkage) of the subset variables selected by DIABLO across the first two latent components. These variables correspond to the molecular features with the highest discriminative power and that contribute to the integration of transcriptomic (black), metabolomic (dark grey), and lipidomic (light grey) datasets. Rows represent selected variables and columns represent samples. Values correspond to standardised expression (z-scores). **C**: Multi-omics relevance network showing correlations between selected transcripts (blue nodes), metabolites (green nodes) and lipidomic families (red nodes). Positive and negative correlations are represented by green and red edges, respectively. Only correlations above the selected threshold (r > 0.7) are displayed. The table summarises the correspondence between the NMR buckets and their associated chemical shift ranges (ppm). Metabolite identification was based on bucket positions, with several buckets potentially corresponding to the same metabolite. **D**: Correlation plots across three omics data based on the first component of DIABLO, displaying strong positive correlations between transcriptome and metabolome (r = 1), transcriptome and lipidome (r = 0.99), as well as metabolome and lipidome (r = 0.99).

Notably, the distance separating planktonic and spatialised populations was greater than that observed between conditions without or with NaCl supplementation. Both spatialised conditions (ST and SNaCl) clustered separately from the planktonic conditions (PT and PNaCl), indicating that growth mode represented the main axis of multi-omics variation under the conditions tested, whereas the contribution of osmotic stress was comparatively smaller. NaCl exposure generated a secondary axis of variation whose magnitude and direction differed between planktonic and spatialised populations, indicating that the response to osmotic stress depended on growth mode. To identify the biological processes underlying this separation, the individual contribution of each variable to global PCA construction was analysed (**Supplementary Dataset 1**). The variables contributing most strongly to PC1 included genes involved in respiratory and energy metabolism (*icdA*, *nuoLMN*, *hycB*), envelope biogenesis and membrane remodelling (*pagO*, *ompW*, *ompX*, *rfa* genes), redox homeostasis (*sel* genes, *suf* operon, *scs* genes) and stress adaptation (*recB, recN, recǪ, rep*), together with several lipidomic (cyclic fatty acids) and metabolomic markers. In contrast, PC2 was mainly driven by variables associated with stress-response pathways, including motility functions (*ffiB*, *ffiC*, *ffgE*), chemotaxis regulators (*cheR* and *cheZ*), and regulatory processes (*greA*, *infB*) involved in environmental adaptation. Overall, this integrative analysis identified spatial organisation as the largest source of multi-omics variation under the conditions tested, with osmotic stress acting on a physiological state already extensively reshaped by spatial growth.

The integrative approach also provided further insight: the clustered image map (CIM) generated from the selected variables showed strong correlations among the omics datasets (**Figure 1B**). Samples were grouped by growth condition, while selected transcriptomic, metabolomic, and lipidomic variables displayed patterns across environments. Therefore, distinct molecular signatures characterised planktonic and spatialised populations, irrespective of the omics layer considered. Network analysis revealed two major multi-omic modules composed of transcriptomic and metabolomic variables, which were linked by a central cluster (**Figure 1C**). The first module was centred on the genes *idnD*, *ilvD, dniR, asrC,* and *rstB*, which were strongly associated with metabolites involved in central carbon metabolism, including formic acid and lactic acid. The second module comprised *STM2374, yaaA, pgpB, STM2800, yihA*, *glyY,* and associated metabolite buckets, suggesting an alternative physiological state. The presence of intermediate nodes connecting both modules suggested a coordinated physiological process rather than independent responses. Strong correlations were observed between the latent components derived from each omics dataset (**Figure 1D**). The first component of the transcriptome was highly correlated with the metabolome (r = 0.99) and lipidome (r = 0.98), and the metabolomic and lipidomic components were also strongly associated (r = 0.97). These results showed that the discriminant signatures extracted from the transcriptomic, metabolomic and lipidomic datasets were strongly associated across the experimental conditions.

### 2. Spatial structuring establishes a biofilm-associated and stress-adaptive transcriptional programme in *Salmonella*

To identify the molecular mechanisms underlying the physiological separation observed in the integrated PCA, transcriptomic profiles of spatialised and planktonic populations, without salt supplementation, were compared. The comparison uses the log2 foldchange (log2FC) average across all conditions as the reference. Differentially Expressed Genes (DEGs) were assigned to Clusters of Orthologous Groups (COG), a functional classification system that groups genes according to their predicted biological roles and cellular processes. This functional classification revealed extensive transcriptional reprogramming associated with spatial structuring (**Figure 2**). Genes upregulated in the spatialised condition (ST) were mainly assigned to ribosome structure and biogenesis (78%), lipid transport and metabolism (75%), secondary metabolites (74%), and inorganic ions and nucleotides transport (64%), whereas downregulated genes involved in motility were predominantly repressed (**Figure 2A**). This functional redistribution has indicated a broad reorganisation of cellular functions during spatial growth, including reduced motility and changes in nutrient acquisition, lipid metabolism, and stress-associated pathways.

**Figure 2:**
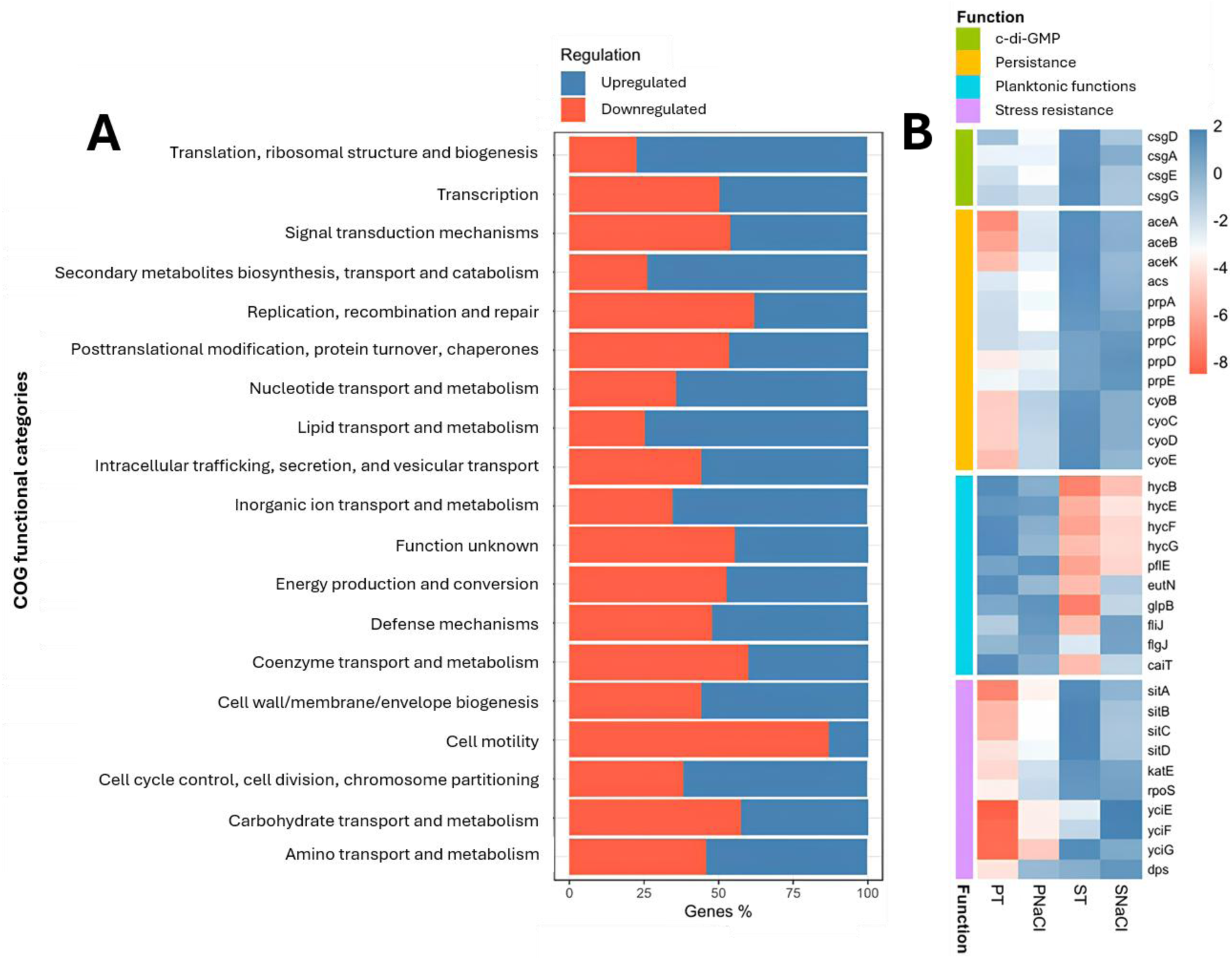
Spatial structuring impacts the distribution of functional categories in *Salmonella* Typhimurium LT2. **A**: Histogram of the proportion of upregulated and downregulated genes across Cluster of Orthologous (COG) categories in the ST vs PT comparison. Downregulated genes were defined by a log2 fold change (log2FC) < 0, whereas upregulated genes were defined by a log2FC > 0. Only genes with an adjusted *p-*value < 0.01 were considered significantly differentially expressed. **B**: Heatmap of representative genes involved in biofilm formation, as known c-di-GMP signalling, persistence, downregulated planktonic functions, and stress resistance. Colours indicate relative expression levels (log2FC) across the planktonic populations without NaCl (PT), planktonic populations supplemented with NaCl (PNaCl), spatialised populations without NaCl supplementation (ST), and spatialised populations supplemented with NaCl (SNaCl). The comparison uses the log2 foldchange (log2FC) average across all conditions as the reference.

To further characterise these lifestyle differences, representative genes involved in biofilm formation, persistence-associated metabolism, stress adaptation, and planktonic functions were examined (**Figure 2B**). Based on the PCA results, lifestyle was the main source of transcriptional variation, whereas osmotic stress had a smaller effect. That is why the expression patterns of these genes were first examined between PT and ST conditions without supplementation with NaCl. The latter responses under osmotic stress conditions (PNaCl and SNaCl) were then considered to determine whether the observed transcriptional regulations were maintained in the presence of NaCl. Moreover, for genes involved in a biofilm-like lifestyle, similar patterns of upregulation or downregulation were observed in the presence or absence of osmotic stress.

A portion of the genes presented in **Figure 2B** is derived from the volcano plot of the ST vs PT DEGs (**Supplementary Figure S2**) and highlights the magnitude of these transcriptional responses. Importantly, spatial structuring promoted the upregulation of genes involved in extracellular matrix production and multicellular behaviour. The upregulation of *csgD, csgA, csgE* and *csgG* suggested activation of the curli-associated regulatory and biosynthetic network. The upregulation of *csgD* together with curli-associated genes and the diguanylate cyclase gene *yaiC* is consistent with activation of a biofilm-associated regulatory network involving c-di-GMP signalling. However, because intracellular c-di-GMP levels were not measured, activation of this second messenger pathway remains to be experimentally confirmed.

Spatialisation was also associated with upregulation of stress-response and nutrient-acquisition systems. A typical pathway upregulated by nutrient limitation and widely associated with long-term survival and persistence is the glyoxylate shunt, which includes the *aceA, aceB,* and *aceK* genes. These genes are upregulated in ST compared to the PT conditions. Additional upregulation of the propionate utilisation genes (*prpA, prpB, prpC, prpD* and *prpE*) and the cytochrome oxidase complex (*cyoB, cyoC, cyoD* and *cyoE*) indicated a shift towards a respiratory and carbon-conserving metabolism. These two pathway genes are highly differentiated, with a log2FC > 3 in the spatialised lifestyle compared to the planktonic lifestyle.

Other pathways supported the hypothesis that spatial organisation promoted adaptation to local nutrient gradients and physicochemical heterogeneity within the hydrogel matrix. For example, the activation of the manganese and iron transporter *sitABCD* was strongly upregulated in the NaCl-free hydrogel compared to PT, along with oxidative and general stress response genes such as *katE, rpoS,* and *dps.* Several poorly characterised but highly responsive genes (*yciE, yciF* and *yciG*) were likewise strongly upregulated in ST compared to PT, suggesting activation of a broad adaptive response. Together, these changes suggested a transition from a growth-oriented metabolism toward a more resource-efficient physiological state.

Conversely, several functions characteristic of actively growing planktonic populations were downregulated. For instance, genes involved in hydrogen metabolism and fermentative pathways (e.g., *hycB, hycE, hycF,* and *hycG*) were strongly downregulated, with an approximately 6-log2FC decrease in ST compared with the mean across all conditions. Genes involved in pyruvate formate metabolism (*pffE*), ethanolamine utilisation (*eutN*), glycerol metabolism (*glpB*), motility (*ffiJ*: -5.45 log2FC in ST compared with the mean across conditions) and chemotaxis-associated functions were also downregulated. The coordinated downregulation of these pathways suggested reduced investment in planktonic growth and motile behaviour. Overall, these trends were consistent with a physiological strategy favouring survival and stress tolerance rather than maximal proliferation.

Several genes selected for detailed functional analysis belonged to the most significantly regulated loci, supporting their contribution to the physiological transition associated with spatial organisation. Collectively, these results demonstrate that growth within the hydrogel upregulated a coordinated transcriptional program that combines specific characteristics of curli-associated functions, carbon-conserving respiratory metabolism, enhanced stress resistance, and repression of classical planktonic functions. To determine whether this transcriptional reprogramming translated into functional metabolic changes, extracellular metabolomic profiles were subsequently analysed.

### 3. The metabolomic profile supports the shift toward a persistence-associated metabolism

To determine whether the transcriptional reprogramming upregulated by spatialised growth translated into functional changes in metabolite consumption and production, extracellular metabolites were monitored during bacterial growth and analysed by ^1^H NMR. First, to follow metabolite evolution during growth and to explain the results obtained after 4 days of growth, a temporal metabolic profile was generated (**Figures 3A and B**).

**Figure 3:**
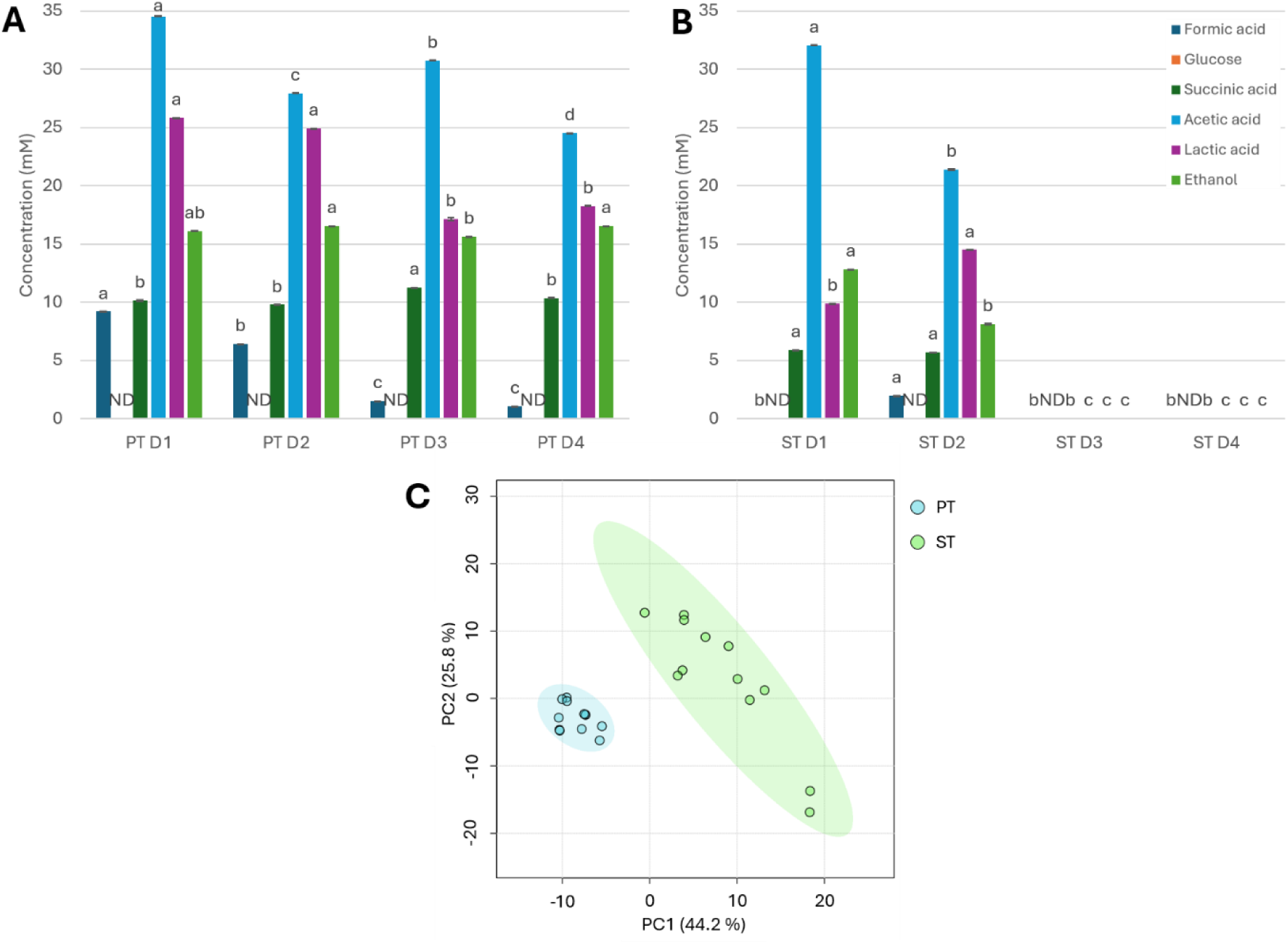
Spatial structuring in hydrogels reshapes the extracellular metabolic profile of *Salmonella* Typhimurium LT2. Kinetics of extracellular metabolites, measured by ^1^H NMR, in **A**: planktonic cultures (PT) or in **B**: spatialised cultures (ST) from day 1 (D1) to day 4 (D4), showing glucose consumption and accumulation of fermentation products, including formic acid, acetic acid, ethanol, lactic acid and succinic acid (n = 3). For each metabolite and condition, differences among incubation times (D1-D4) were assessed using one-way ANOVA followed by Tukey HSD post hoc test (α = 0.05). Different letters denote statistically significant differences between days, whereas groups sharing at least one common letter do not differ significantly (ND: Not Determined). **C**: PCA of the extracellular metabolomic profiles measured at D4 by ^1^H NMR (n = 12). PC1 and PC2 explain 70% of the total variance and discriminate between planktonic and spatialised populations, indicating the establishment of distinct metabolic states.

Kinetics revealed a marked difference between planktonic and spatialised populations. In both conditions, glucose was consumed during the first day of growth, indicating active carbon utilisation. However, the use of glucose carbon varied with lifestyle, as shown by the NMR spectra in **Supplementary Figures S3 and S4**. Planktonic populations (PT) accumulated large amounts of acetic acid (35 mM), lactic acid (25 mM), ethanol (16 mM), succinic acid (10 mM) and formic acid (9 mM) at D1 (**Figure 3A**). These metabolites remained abundant throughout the four-day growth, except for formic acid and lactic acid, which decreased over time, consistent with sustained fermentative activity. In contrast, microcolonies showed lower metabolite concentrations over time, except for acetic acid, despite similar glucose availability (14 mM) and depletion. During the early stages of growth (D1-D2), organic acids and ethanol are accumulated in the hydrogel, indicating that these metabolites were initially produced by the bacteria. However, these concentrations were significantly reduced and became undetectable at D3-D4 (**Figure 3B**) (limit of detection between 0.1 and 1 mM), indicating a second phase of metabolite consumption. Experiments carried out on PT and ST media without *Salmonella,* but spiked with the detected metabolites, showed that metabolite concentrations remained unchanged over time (**Supplementary Figure S5**). Therefore, the disappearance of metabolites observed in ST could not be attributed to evaporation, spontaneous degradation or interactions with the hydrogel matrix, but rather reflected active bacterial uptake and metabolism. This pattern suggests that cells organised in microcolonies not only produce metabolic products but also reuse them during growth. These observations indicate greater net accumulation of classical fermentation end-products in planktonic cultures, whereas their reduced accumulation and subsequent disappearance in spatialised cultures are consistent with altered production and/or reutilisation.

Multivariate analysis further supported the existence of distinct metabolic states. PCA performed on metabolomic profiles collected at D4 (same time as transcriptomic analysis) showed a clear separation between planktonic and spatialised populations along PC1, which explained 44.2 % of the total variance (**Figure 3C**).

These temporal changes were consistent with the previously analysed transcriptomic data. Several genes associated with fermentative and anaerobic metabolism, including members of the *hyc* operon and *pffE*, were downregulated in spatialised cells (**Figure 2B**), consistent with reduced accumulation and depletion of formic acid. For this reason, the reduced accumulation and subsequent consumption of formic acid, ethanol and lactic acid supported a decreased reliance on fermentative metabolism. Conversely, genes involved in aerobic respiration, including the cytochrome bo_3_ oxidase complex (*cyoB, cyoC, cyoD,* and *cyoE*), were upregulated in microcolonies, suggesting an increased reliance on aerobic respiratory pathways.

Concerning the central carbon pathways, a substantial remodelling was associated with the spatialisation of *Salmonella* cells. Spatialised populations displayed an upregulation of the *acs* gene, which encodes acetyl-CoA synthetase. This transcriptional change was accompanied by a marked decrease in extracellular acetate concentrations during the later stages of growth, indicating acetate reutilisation. Moreover, the strong activation of the glyoxylate shunt genes *aceA, aceB,* and *aceK*, the propionate utilisation pathway (*prp* operon), and several C4-dicarboxylate transport systems indicated extensive remodelling of succinic acid-associated carbon fluxes. In addition, the induction of *dld* by microcolonies, encoding a D-lactate dehydrogenase, coincided with the progressive disappearance of lactate in ST, supporting the active reutilisation of this organic acid. Taken together, these data indicate that spatialisation is associated with a profound reorganisation of carbon fluxes involving the glyoxylate shunt, fermentative metabolism and respiratory pathways. Accordingly, this metabolic shift translated into a more resource-efficient and biofilm-associated mode of carbon utilisation. The extracellular metabolomic profiles, therefore, provide functional support for the metabolic reprogramming inferred from the transcriptomic data.

### 4. Spatial structuring induces extensive remodelling of the bacterial membrane

To determine whether the transcriptomic and metabolic reprogramming upregulated by spatial organisation extended to membrane architecture, lipid-associated pathways were analysed at both the transcriptomic and lipidomic levels in parallel. First, differentially expressed genes (DEGs), identified between ST and PT by RNA-seq and assigned to the COG category of lipid transport and metabolism, were mapped to functional pathways using KEGG annotations (**Figure 4A**). In parallel, membrane fatty acid (FA) composition was characterised by gas chromatography coupled with mass spectrometry (GC-MS) and classified into four major FA families: saturated fatty acids (SFA), unsaturated fatty acids (UFA), cyclopropane fatty acids (cycloFA) and hydroxylated fatty acids (HFA) (**Figure 4B**). To identify the molecular basis underlying these changes, DEGs were mapped onto FA and glycerophospholipid metabolic pathways, using the KEGG platform (**Figure 4C**). Until now, our analysis had mainly focused on the global impact of spatial structuring on transcriptomic and metabolomic profiles. In this section, we examined whether exposure to osmotic food-related stress (6% NaCl) also influenced the omics response, with particularly marked effects on membrane lipid composition.

**Figure 4:**
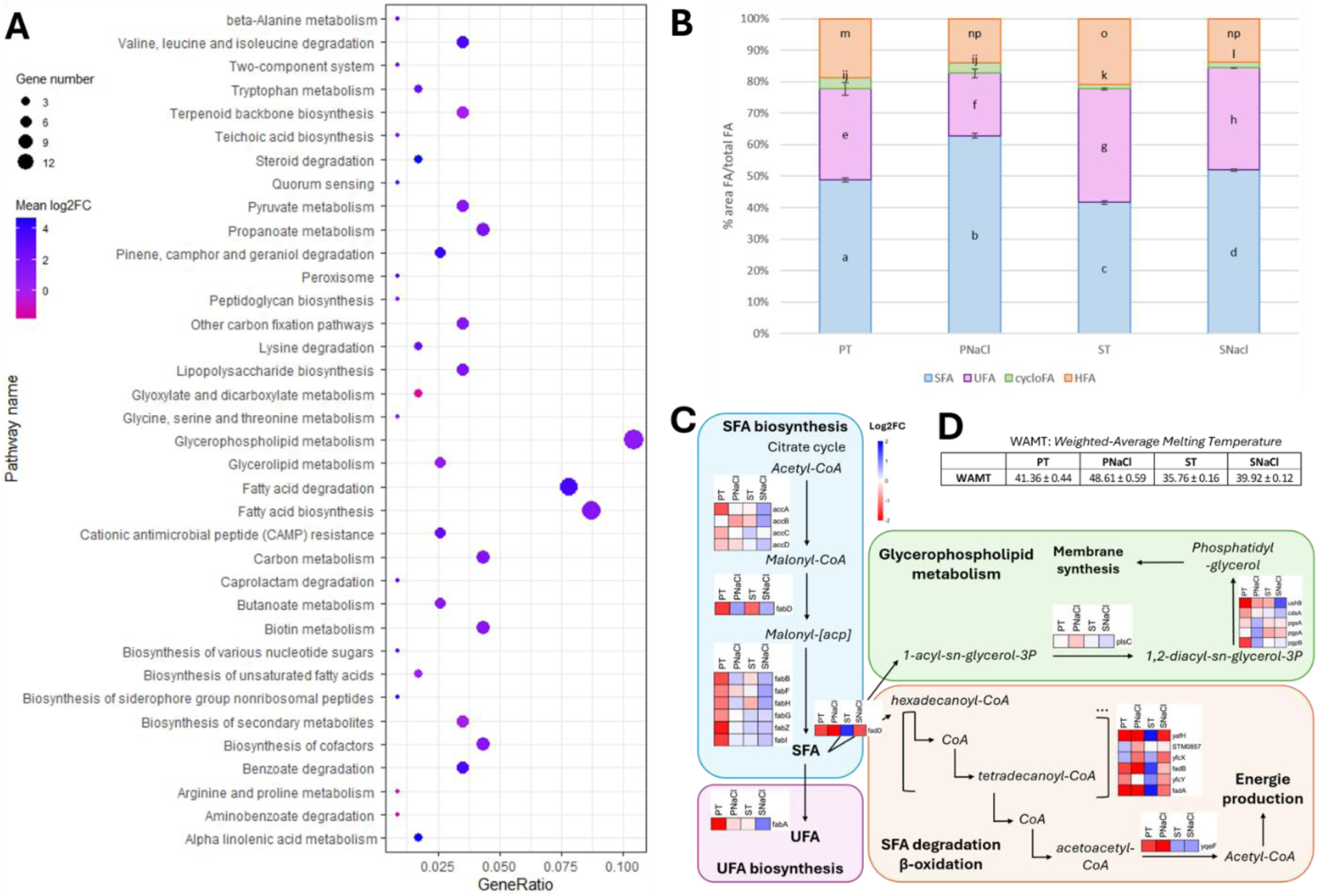
Spatial structuring reshapes membrane lipid metabolism and composition in *Salmonella* Typhimurium LT2. **A**: Functional enrichment by KEGG (Kyoto Encyclopedia of Genes and Genomes) of lipid transport and metabolism GOG (Cluster of Orthologous Groups) pathway (n = 3). Bubble size represents the number of genes (*p*_adj_ < 0.01) assigned to each pathway, and colour indicates the mean log2FC from the ST vs PT DEGs. Results were obtained from RNA-seq analysis. **B**: Relative distribution of major fatty-acid classes in planktonic NaCl-free (PT), planktonic supplemented with 6% NaCl (PNaCl), spatialised NaCl-free (ST) and spatialised supplemented with 6% NaCl (SNaCl) environment obtained by GC-MS analysis. Values are presented as mean ± standard deviation (SD) (n = 9). Statistical significance was assessed by one-way ANOVA followed by Tukey’s HSD test. Means that do not share a common letter are significantly different (p < 0.05). SFA: Saturated Fatty Acids; UFA: Unsaturated Fatty Acids; cycloFA: cyclopropane Fatty Acids; HFA: Hydroxylated Fatty Acids. **C**: Heatmaps of transcriptomic changes affecting pathways of fatty-acid biosynthesis, β-oxidation and glycerophospholipid metabolism (n = 3). Colour scale indicates relative gene expression changes taken as reference the log2FC average of all conditions. Scale has been normalised between -2 and 2 log2FC. **D**: Weighted-average melting temperature (WAMT) values for each *S.* Typhimurium growth condition. WAMT values indicate bacterial membrane fluidity: the higher the WAMT, the more rigid the membrane; conversely, a lower WAMT indicates greater membrane fluidity.

Functional enrichment analysis revealed that spatialisation (ST vs PT) affected multiple pathways involved in lipid transport and metabolism within the COG category (**Figure 4A**). Among the most enriched pathways were glycerophospholipid metabolism (11%), FA biosynthesis (8.7%) and FA degradation (7.6%). Several central carbon metabolic pathways that feed into lipid synthesis, including glyoxylate and dicarboxylate metabolism (2%) and propanoate metabolism (4%), were also significantly enriched. These pathways connect acetyl-CoA metabolism, carbon reutilisation and the production of biosynthetic intermediates (∼25% of the gene ratio), suggesting extensive coupling between carbon metabolism and membrane remodelling.

Lipidomic analysis by GC-MS confirmed that spatialisation profoundly altered membrane composition (**Figure 4B**). Compared with planktonic populations, spatialised cells displayed a significantly lower relative abundance of SFA and a concomitant increase in UFA. Exposure to 6% NaCl upregulated additional modifications in FA composition, although the magnitude of these changes depended on the bacterial lifestyle. In planktonic populations, NaCl promoted a marked increase in the relative abundance of SFA together with a reduction in UFA, consistent with membrane rigidification. By contrast, spatialised populations displayed more limited changes following salt exposure, with the overall lipid profile remaining closer to that observed under NaCl-free spatialised conditions. Together, these results demonstrated that both spatialisation and osmotic stress alter membrane composition, but that spatialisation is the dominant factor structuring the lipidome, in agreement with the global multi-omics analyses presented above.

Changes were also observed in the HFA and cycloFA fractions, but given their low abundance, these changes should be interpreted with caution, as they may primarily reflect shifts in relative abundance (% of the total pool) rather than significant changes in absolute concentration. HFA, which are components of the outer membrane of Gram-negative bacteria, are synthesised through FA hydroxylation mediated by the *lpxO* gene. As *lpxO* expression remained unchanged across the four growth conditions, correlated with no significant differences in HFA levels. Likewise, cycloFA are generated either *de novo* or by post-synthetic conversion catalysed by *cfa*. The stable expression of *cfa* across conditions was consistent with the absence of significant changes in cycloFA abundance.

Spatialisation affected all major stages of membrane lipid metabolism, including FA biosynthesis, FA degradation, and membrane glycerophospholipid synthesis. As shown in the heatmaps, numerous genes were differentially expressed (**Figure 4C**). Compared with planktonic conditions without NaCl, genes involved in SFA biosynthesis were upregulated, particularly the *acc* and *fab* operons. The first catalyses the conversion of acetyl-CoA to malonyl-CoA, and the second catalyses the elongation and modification of acyl chains, leading to the synthesis of SFA. UFAs are synthesised from SFAs by the *fabA* gene, whose expression is downregulated in PT compared to ST. The presence of NaCl in the medium increased the expression of *fabA* in both lifestyles. In contrast, the regulation of *fadD* indicates substantial remodelling of β-oxidation and glycerophospholipid metabolism. The latter gene was overexpressed in ST compared to PT. Nevertheless, in the presence of 6% NaCl, genes associated with both pathways were downregulated in planktonic and spatialised cultures. Continuing along these pathways, genes involved in SFA degradation were regulated similarly: *fadD* showed reduced expression in the planktonic lifestyle and in SNaCl, whereas it was globally upregulated in ST compared with the mean expression level across all conditions. In addition, several genes involved in glycerophospholipid synthesis, including *gpsA, plsC*, and *ushB* were up regulated.

To assess the functional consequences of these lipidomic changes on membrane physicochemical properties, the weighted average melting temperature (WAMT) was calculated for each condition (**Figure 4D**). WAMT is commonly used as an indicator of membrane rigidity, with higher values corresponding to less fluid membranes. Spatialisation alone resulted in a decrease in WAMT, from 41.36 ± 0.44 °C in PT to 35.76 ± 0.16 °C in ST, indicating that growth in microcolonies maintained more fluid membranes than planktonic cells. In contrast, exposure to 6% NaCl increased WAMT values in both lifestyles, reaching 48.61 ± 0.59 °C in PNaCl and 39.92 ± 0.12 °C in SNaCl, consistent with membrane rigidification under osmotic stress. Despite this increase, the membrane of cells organised in microcolonies and exposed to NaCl remained substantially more fluid (+ 4.16 °C) than that of PNaCl populations (+ 7.25 °C). These results demonstrate that spatial structuring establishes a distinct membrane state characterised by increased fluidity, which is only partially counterbalanced by osmotic adaptation.

Finally, spatialised populations were characterised by higher expression of the β-oxidation pathway, concomitant with lower SFA concentrations. Together, the KEGG enrichment analysis (**Figure 4A**), the FA composition (**Figure 4B**), the coordinated regulation of lipid metabolic pathways (**Figure 4C**), and the final membrane fluidity calculation (**Figure 4D**), demonstrated that spatial structuring upregulated large-scale remodelling of the bacterial membrane. Because membrane composition directly influences permeability and osmotic adaptation, these structural modifications may have contributed to the distinct responses between planktonic and spatialised populations in high-salt conditions.

### 5. Spatial structuring leads to extensive reprogramming of the osmotic stress response

To determine whether the physiological state established by spatial structuring influenced adaptation to osmotic stress, transcriptomic and metabolomic responses to 6% NaCl were compared between planktonic and spatialised populations (**Figure 5**). DEGs were analysed by RNA-seq and assigned to COG functional categories (**Figures 5A and B**), while extracellular metabolic profiles were characterised by ^1^H NMR (**Figures 5C and D**).

**Figure 5:**
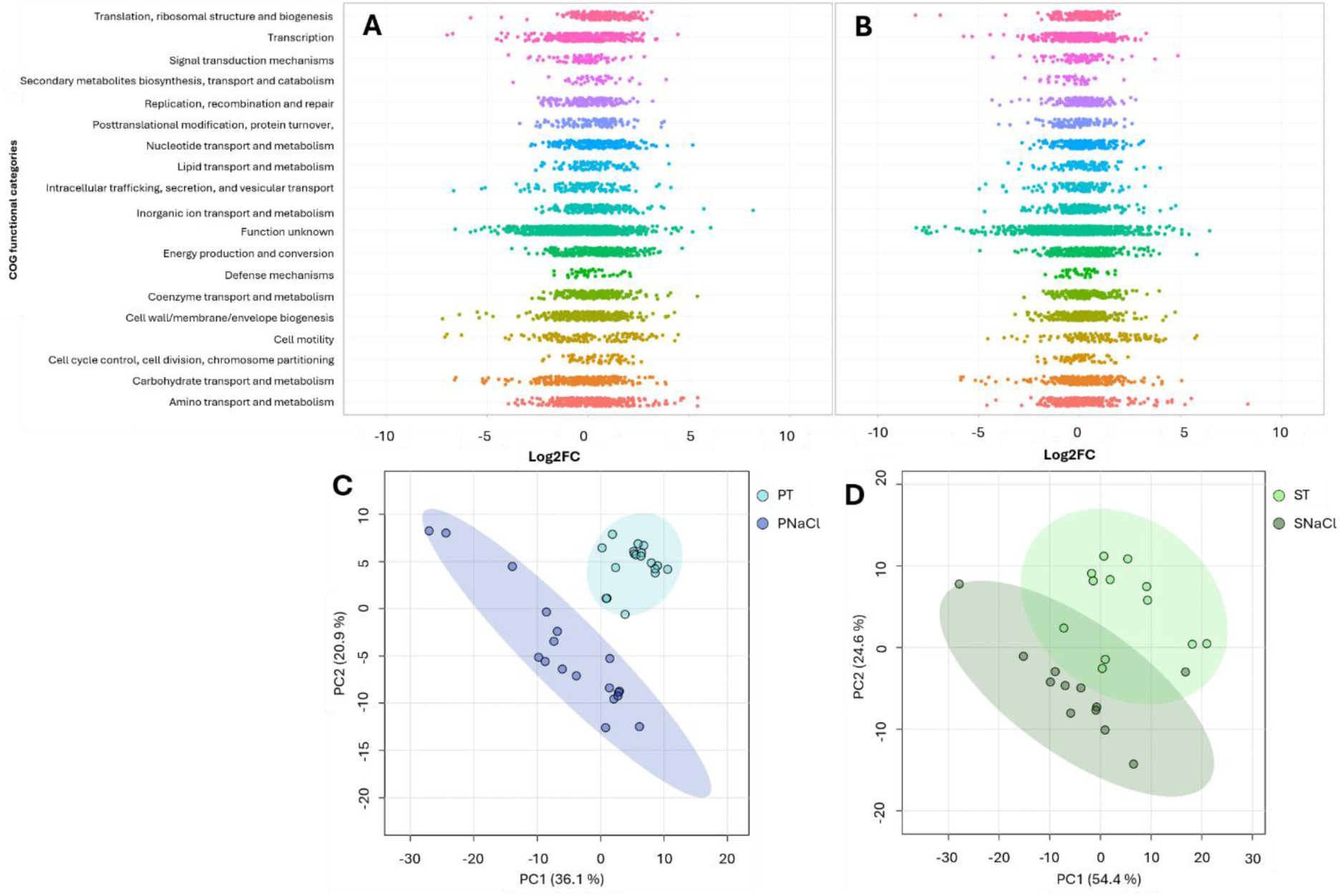
Spatial structuring reprograms transcriptional and metabolic responses to osmotic stress in *Salmonella* Typhimurium LT2 bacteria. Distribution of DEG across COG functional categories in populations exposed to 6% NaCl. Each point represents the log2FC value of a gene obtained from DEG between **A**: PNaCl vs PT (n = 18) or **B**: SNaCl vs ST (n = 12). PCA of extracellular metabolomic profiles at D4 analysed by ^1^H NMR, of **C**: planktonic or **D**: spatialised populations exposed or not to 6% NaCl.

Functional classification of DEGs revealed distinct transcriptional responses to NaCl across bacterial lifestyles. For the two DEG analyses, quantitative data (mean log2FC for each COG) were provided in **Supplementary Dataset 2**. In planktonic populations, osmotic stress was associated with broad transcriptional changes affecting numerous COG categories (**Figure 5A**). In particular, the translation and ribosomal biogenesis pathway (mean log2FC = 0.72), energy production and conversion pathway (mean log2FC = 0.43), cell cycle control, cell division, chromosome partitioning pathway (mean log2FC = 0.42) and nucleotide transport and metabolism pathway (mean log2FC = 0.41) were upregulated in planktonic populations exposed to osmotic stress compared to growth in the PT populations. In contrast, spatialised populations generally displayed lower average transcriptional shifts, except for the cell motility pathway, which is strongly upregulated in the presence of 6% NaCl (meanlog2FC = 1.03) (**Figure 5B**). Secondary metabolite biosynthesis, transport, and catabolism pathways were downregulated in spatially isolated populations exposed to osmotic stress (mean log2FC = -0.37). These observations indicated a globally attenuated transcriptional response of microcolonies to osmotic stress.

A common transcriptional response was nevertheless detected in both lifestyles. This shared response included upregulation of compatible solute transport systems, particularly the *proVWX* operon, along with genes involved in polyamine metabolism and flagellar assembly. These changes revealed a conserved core response to osmotic stress that was independent of growth mode.

The extracellular metabolomic profiles were consistent with these observations. PCA of extracellular metabolite profiles at D4 revealed a clear separation between PT and PNaCl populations (PC1 = 36.1 %; PC2 = 20.9 % of the total variance), indicating major metabolic reprogramming following NaCl exposure in planktonic cells (**Figure 5C**). Quantitative analysis of the identified metabolites showed that group separation was associated with changes in organic acids and fermentation-related compounds, including lactate, formate, acetate, succinate, and ethanol. In planktonic populations, NaCl altered the accumulation of these extracellular metabolites, likely due to differences in growth-rate kinetics, particularly an extended Lag phase under osmotic stress (**Hoch et al. 2026**). Osmotic stress also altered the metabolomic profile of spatially isolated populations (**Figure 5D**). PC1 and PC2 explained 54.4% and 21.6% of the variance, respectively. ST and SNaCl samples occupied different regions of the PCA space, although their confidence ellipses partially overlapped. Moreover, greater biological variability was observed in microcolonies than in planktonic populations.

Monitoring metabolite dynamics over time provided further indication of the differences observed between the growth conditions (without and with 6% NaCl) and between the two lifestyles. The metabolite quantification is presented in **Figures 3A and B** for NaCl-free conditions and in **Supplementary Figure S6** for osmotic stress. For growth under planktonic conditions, the major difference between the NaCl-free and osmotic stress was the metabolite levels quantified after four growing days, notably for lactic and acetic acid (higher in the presence of NaCl). In contrast, in the hydrogel, the presence of 6% NaCl prevented the complete disappearance of several metabolites that were observed in ST at D4. The more likely reason for this difference may be the state of growth kinetics, as the presence of osmotic stress extended the lag phase.

The transcriptomic and metabolomic datasets converged towards the same conclusion. Planktonic populations exhibited extensive transcriptional and metabolic reorganisation in response to NaCl, whereas spatialised populations showed a more limited shift from their pre-existing physiological state. These pathway reorganisations are illustrated in **Table 1**.

**Table 1:**
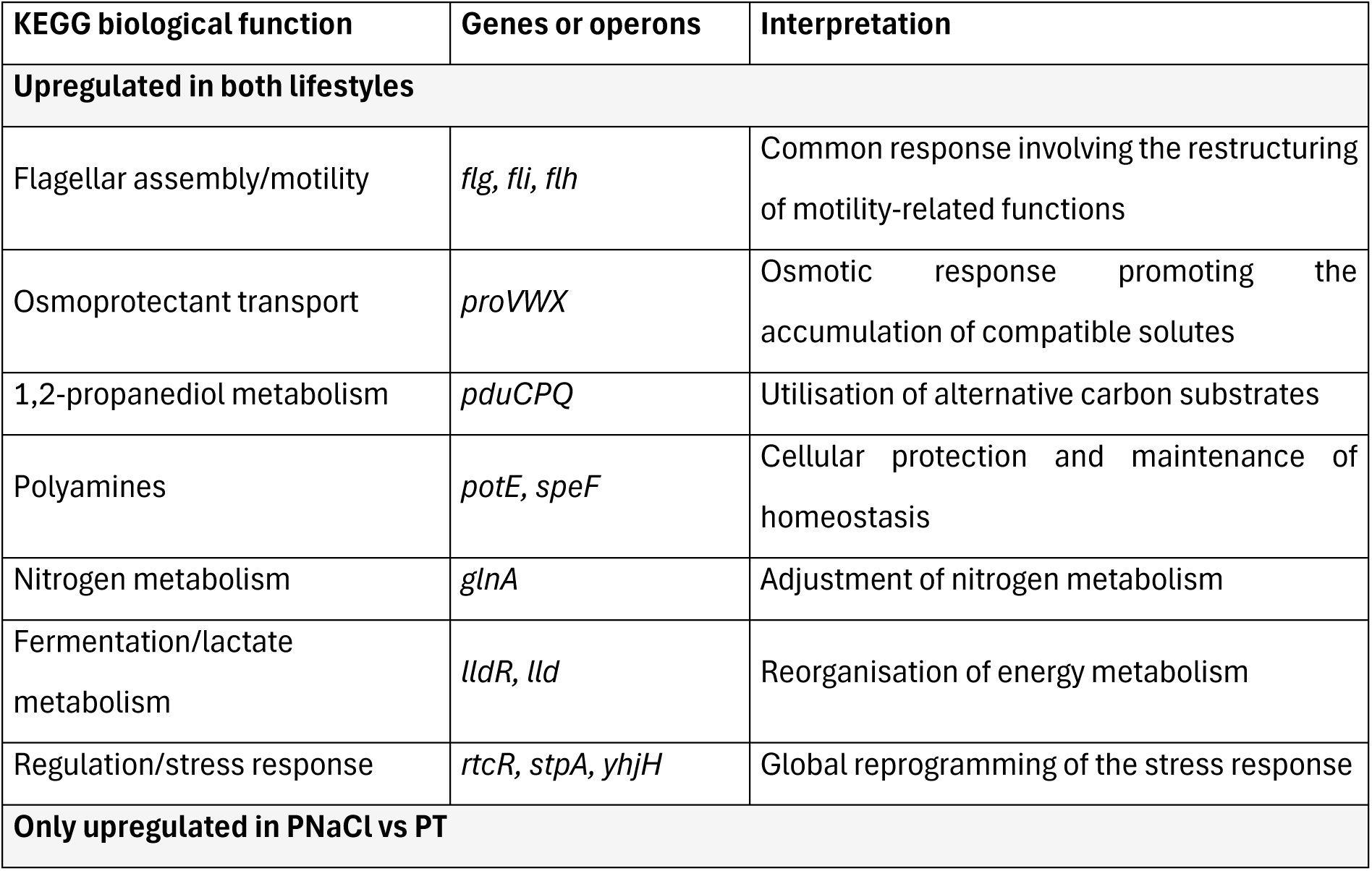
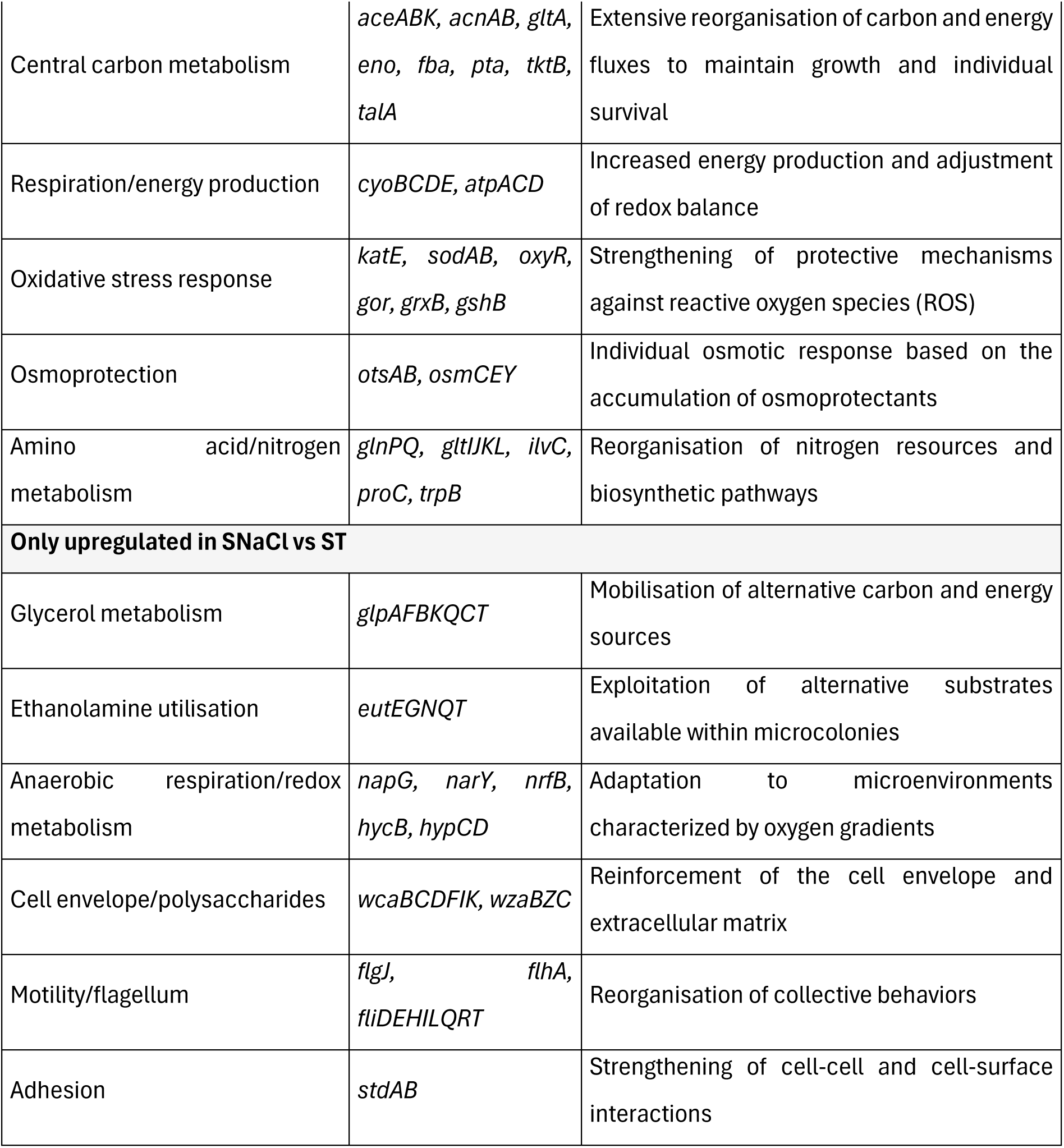
Functional classification of major transcriptional responses to osmotic stress in planktonic and spatialised populations of *Salmonella* Typhimurium LT2. Genes were grouped according to their major KEGG-associated biological functions. Only representative genes belonging to pathways relevant to osmotic adaptation are shown. Genes upregulated in both lifestyles correspond to log2FC > 2 in both PNaCl vs PT and SNaCl vs ST comparisons. Lifestyle-specific genes correspond to log2FC > 2 in only one comparison. The complete list of DEGs and associated KEGG pathways is provided in **Supplementary Dataset 3**.

**Table 1:**
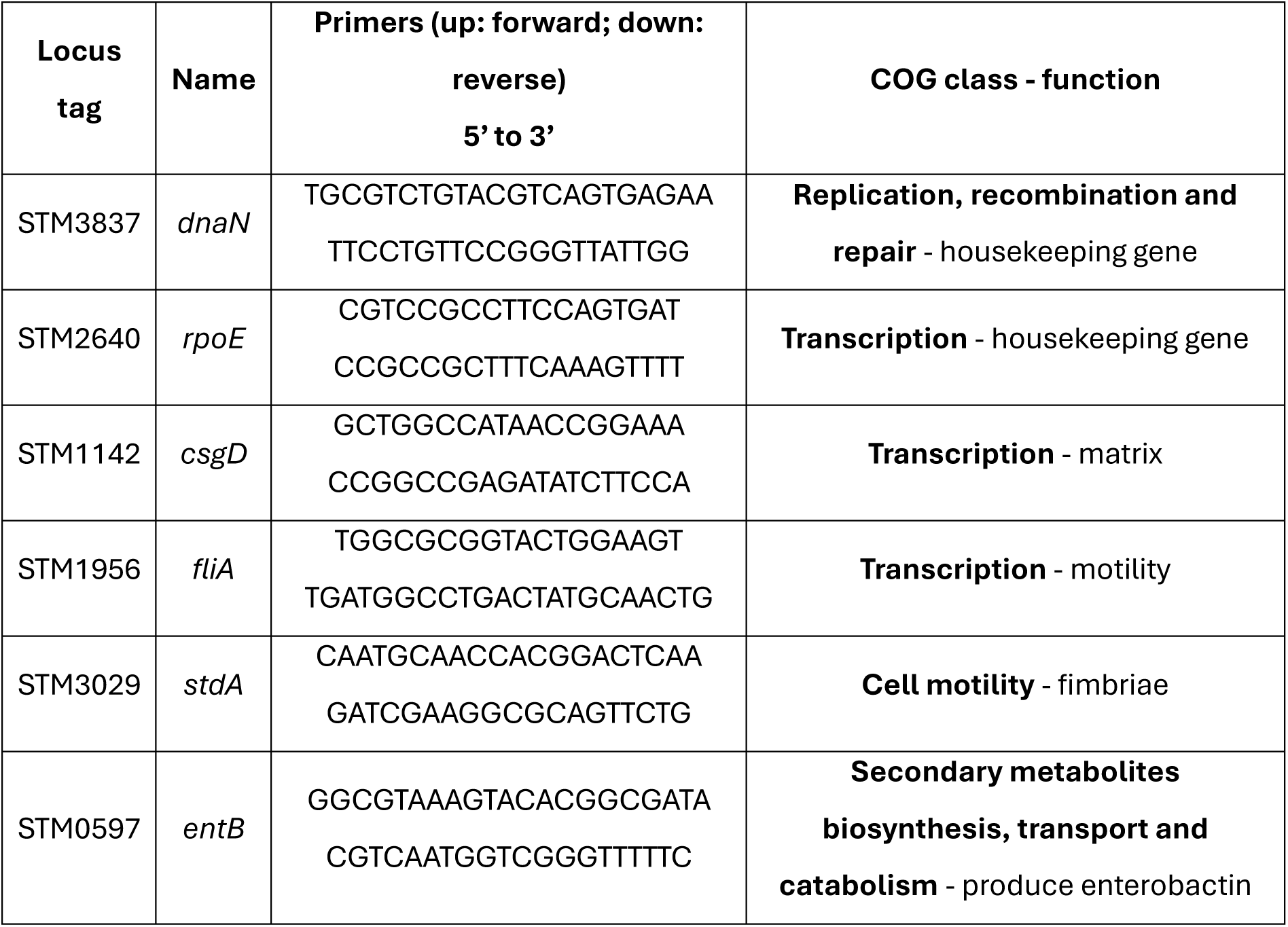
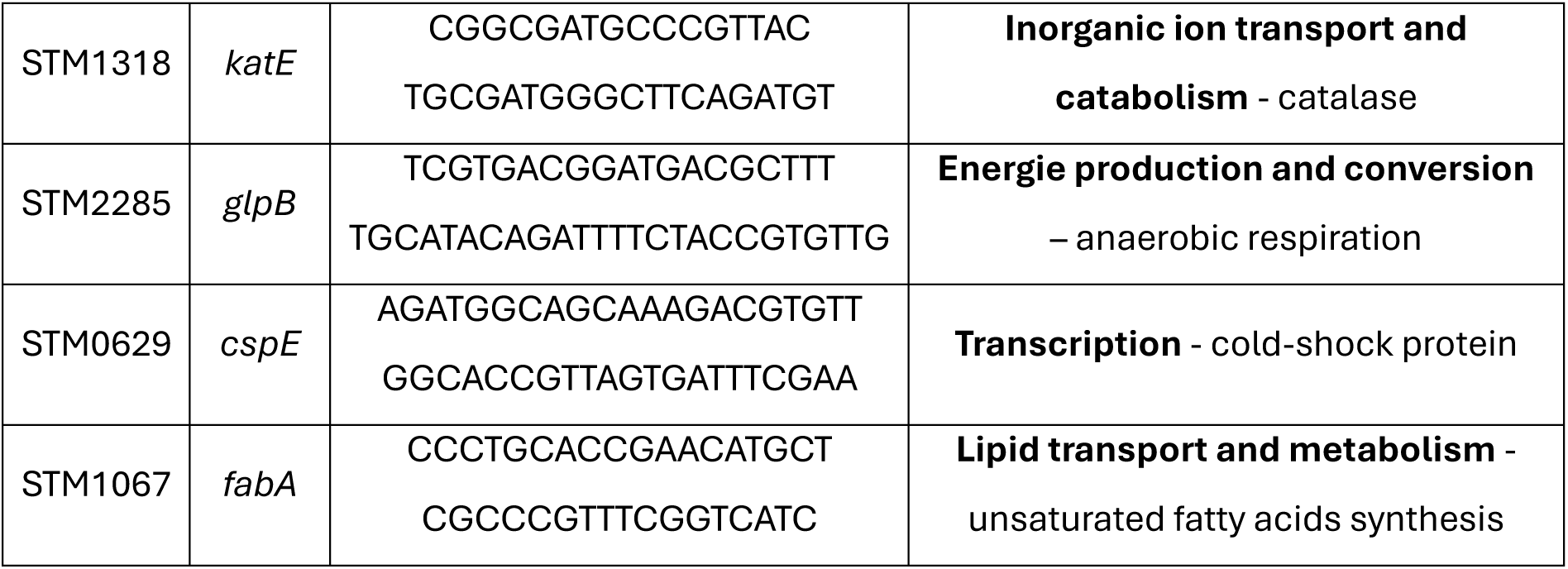
*Salmonella* Typhimurium LT2 genes targeted in the RT-qPCR analysis classified by function.

In the presence of 6% NaCl in the environment, both planktonic and spatialised populations shared a core response major involved in osmotic homeostasis (*proVWX*) and motility-associated function (*ffg* and *ffi* operons). As noted above, NaCl was associated with the upregulation of motility related genes, in both free-living cells and cells organised into microcolonies within a hydrogel matrix. To further investigate the effects of osmotic stress on motility-related phenotypes, confocal microscopy analyses were performed. The results are presented in the **Supplementary Figure S7**.

Beyond this common adaptive program, planktonic populations preferentially upregulated pathways associated with carbon metabolism (*aceABK*, *acnAB*, etc.), energy production (*cyoBCDE*), oxidative stress protection and osmoprotection. These functions were accompanied by the strong metabolic reorganisation observed in the NMR data and are consistent with the extensive transcriptional response detected in PNaCl populations. In contrast, spatialised populations preferentially upregulated pathways related to polysaccharides biosynthesis (*wcaBCDFIK*), adhesion (*stdAB*), alternative carbon utilisation (*glpAFBKǪCT*) and anaerobic respiration. The enrichment of these functions suggested that adaptation to osmotic stress in bacteria organised as microcolonies relied not only on cellular osmoprotection but also on processes potentially contributing to collective protection and exploitation of heterogeneous microenvironments.

Together, these results suggest that the physiological state established during spatial growth modifies the subsequent response to osmotic stress, reducing the extent of transcriptional and metabolic reorganisation observed after salt exposure. Nevertheless, does this collective protection represents a uniform response, or is there local heterogeneity at the scale of the microcolony?

### 6. Targeted spatial gene-expression analysis reveals microscale heterogeneity within *Salmonella* microcolonies

To investigate how these key adaptive functions are distributed within spatially organised populations, a spatial transcriptomic approach was implemented at the microcolony scale. Microcolonies were first chemically fixed to enable downstream processing on the @Bridge platform (INRAE, Jouy-en-Josas). The central and peripheral regions of individual microcolonies were subsequently isolated by laser capture microdissection (LCM), with calcofluor white staining to delineate colony structure and guide region collection (see **Supplementary Figure S8**). Total RNA was extracted independently from each region, amplified, and analysed by RT-qPCR using a panel of genes associated with biofilm formation (*csgD*), stress adaptation (*cspE, katE*), motility (*ffiA*), membrane remodelling (*fabA*), iron acquisition (*entB*), adhesion (*stdA*) and carbon metabolism (*glpB*). Preliminary validation experiments confirmed the linearity of RNA amplification over the range of RNA concentrations recovered following LCM (see **Supplementary Figure SG**). A constant ΔCt across the different genes was considered indicative of comparable amplification efficiency for all target genes. Threshold cycle (Ct) values obtained from RT-qPCR were then used to generate the spatial transcriptional profiles presented below. Two housekeeping genes were used to normalise the gene expression: *dnaN* and *rpoE*. The Ct variability is presented in **Supplementary Figure S10,** with coefficients of variation of 6.75 for *dnaN* and 4.77 for *rpoE*.

PCA revealed clear structuring of transcriptional profiles by both osmotic stress and microcolony location, except for certain biological replicates (**Figure 6**). The first principal component (PC1), which accounted for 71.1% of the total variance, separated microcolonies exposed to NaCl from NaCl-free microcolonies, except for biological replicate 2 of SFTp. Globally, this separation indicated that osmotic stress remained the dominant determinant of gene expression in spatially structured populations, consistent with global RNA-seq results. The second principal component (PC2), explaining an additional 14.5% of the variance, separated central and peripheral regions, except for two different biological replicates, SFNaClp_3 and SFNaClc_2. Nevertheless, these results suggested the presence of a small degree of intra-colony transcriptional heterogeneity.

**Figure 6:**
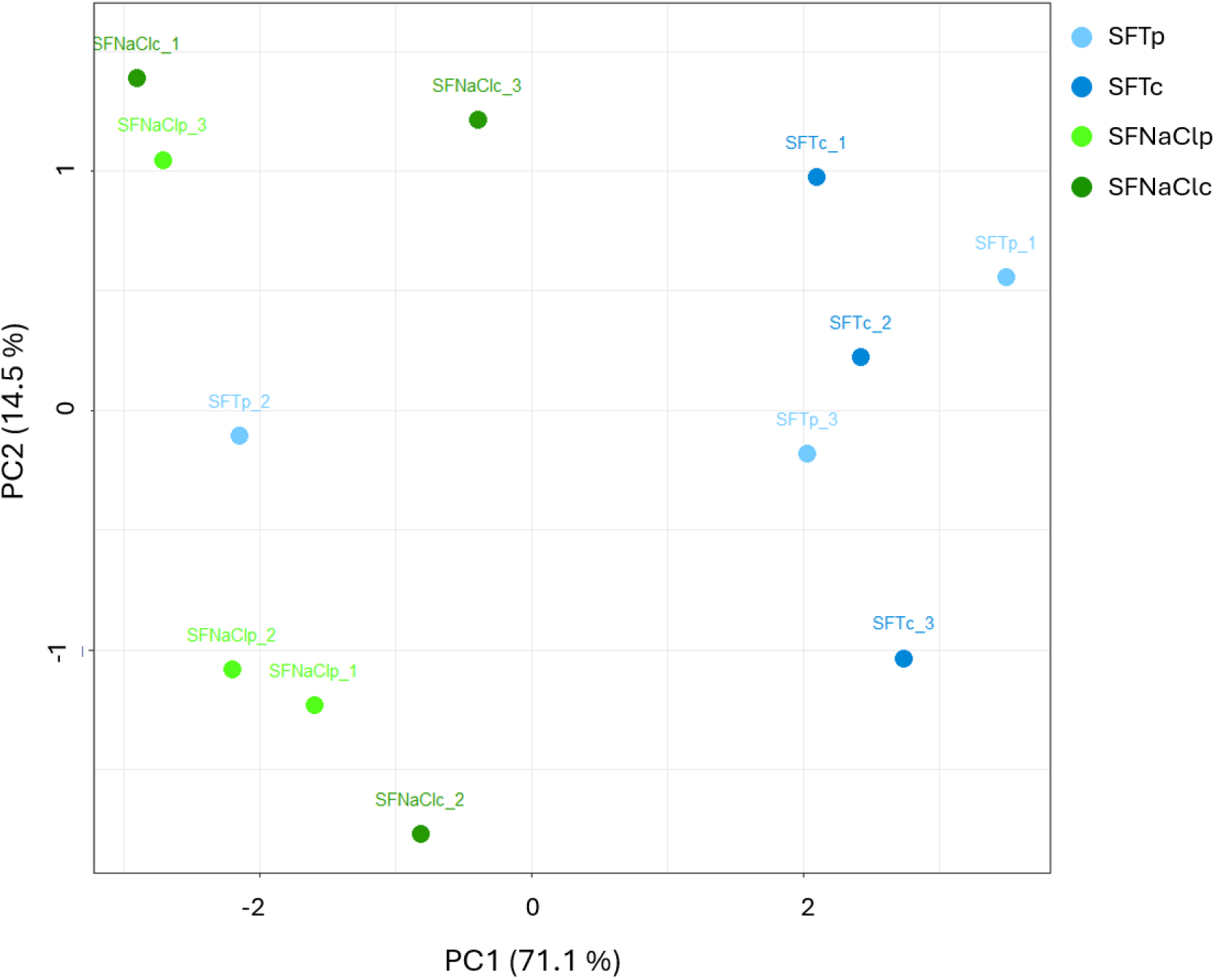
Principal component analysis (PCA) of RT-qPCR expression profiles obtained from the centre (c) and periphery (p) of fixed (F) *Salmonella* Typhimurium LT2 microcolonies grown in the absence (ST) or presence (SNaCl) of 6% NaCl. Expression profiles were generated using *csgD, cspE, katE, ffiA, fabA, entB, stdA* and *glpB* genes. PC1 and PC2 explain, respectively, 71.1% and 14.5% of the total variance.

Interestingly, peripheral and central regions were grouped together yet remained distinct under osmotic stress, suggesting that spatial heterogeneity persists despite the strong global effect of this stress. These results demonstrated that the physiological adaptation identified by population-level transcriptomic analyses was not uniformly distributed across the microcolony, highlighting the need to further study these responses at a more spatially resolved scale.

Consistent with the PCA analysis, the log2FC calculation revealed substantial variation among biological replicates (see **Supplementary Figure S11**). Nevertheless, the average transcriptional responses were consistent with the trends identified in the RNA-seq dataset. In particular, *ffiA* (log2FC =3.27) and *stdA* (log2FC = 2.22) showed a significant difference between the centre of microcolonies grown without or with 6% NaCl. The genes are upregulated in the presence of osmotic stress, concomitant with global RNA-seq analysis (*ffiA* log2FC SNaCl vs ST = 3.03; *stdA* log2FC SNaCl vs ST = 6.01). Similarly, *csgD, entB, fabA,* and *glpB* were upregulated under osmotic stress. In contrast, *cspE,* involved in stress adaptation and virulence, was downregulated under osmotic stress, consistent with RNA-seq analysis, which showed a log2FC of -5.32 in the DEG SNaCl vs ST. Finally, *katE* was the least affected gene by the condition studied here. These results validated the spatial transcriptomic workflow and indicated that the major transcriptional signatures identified at the population level were also detectable at the microcolony scale.

Interestingly, inspection of individual biological replicates revealed spatial expression gradients that were not always apparent when averaging all replicates, because RT-qPCR results are not fully reproducible. For this reason, subsequent analyses focused on transcriptional heterogeneity within a single biological replicate (corresponding to a pool of ∼15-20 microcolonies), rather than on mean expression. Comparison between conditions, and more precisely between the centre and the peripheries of microcolonies, revealed several recurring spatial expression patterns (see **Supplementary Dataset 4**).

Comparison of gene expression between the centre and the periphery of microcolonies revealed several recurring spatial patterns. In the absence of NaCl, genes associated with active growth tended to be preferentially expressed at the colony periphery. This trend was particularly apparent for *stdA*, implicated in adhesion and whose expression was higher in peripheral regions in the three replicates. Similar peripheral enrichment tendencies were observed for *csgD, entB, fabA, ffiA* and *glpB*, although the magnitude of the gradients varied between replicates. Together, these observations suggested that cells at the colony periphery were generally more metabolically active than those in the centre. In contrast, genes associated with stress adaptation displayed the opposite tendency. Gene coding for cold-shock proteins *cspE* (mean log2FC SFTc vs SFTp = 0.22) wwas generally more expressed in the centre of the microcolonies, consistent with the establishment of local microenvironments. These conditions were expected to impose physiological constraints that favour the activation of stress-protection mechanisms within the colony interior. Interestingly, exposure to NaCl did not significantly modify the pattern of gene expression, except for *ffiA*. Osmotic stress was associated with reduced *ffiA* expression (involved in motility) at the microcolony periphery.

To independently examine spatial patterns of promoter activity within microcolonies, fluorescent transcriptional reporters for *ffiA* and *csgD* were analysed in the closely related *S. enterica* serovar Typhimurium ATCC 14028 background (**Figure 7**). ATCC 14028 displayed growth, osmotic adaptation, and gastrointestinal tolerance phenotypes broadly comparable to those of LT2 (**Supplementary Table S1**), providing a phenotypically relevant background for complementary reporter-based analyses. Wild-type strains ATCC 14028 and LT2 were used as negative controls to assess laser penetration throughout the entire microcolony and to ensure the absence of green autofluorescence (**Supplementary Figure S12**). Kymograph analysis showed that the red fluorescent signal from Syto61 was detected throughout the microcolony (approximately 60 µm in diameter), while no green autofluorescence was detected.

**Figure 7:**
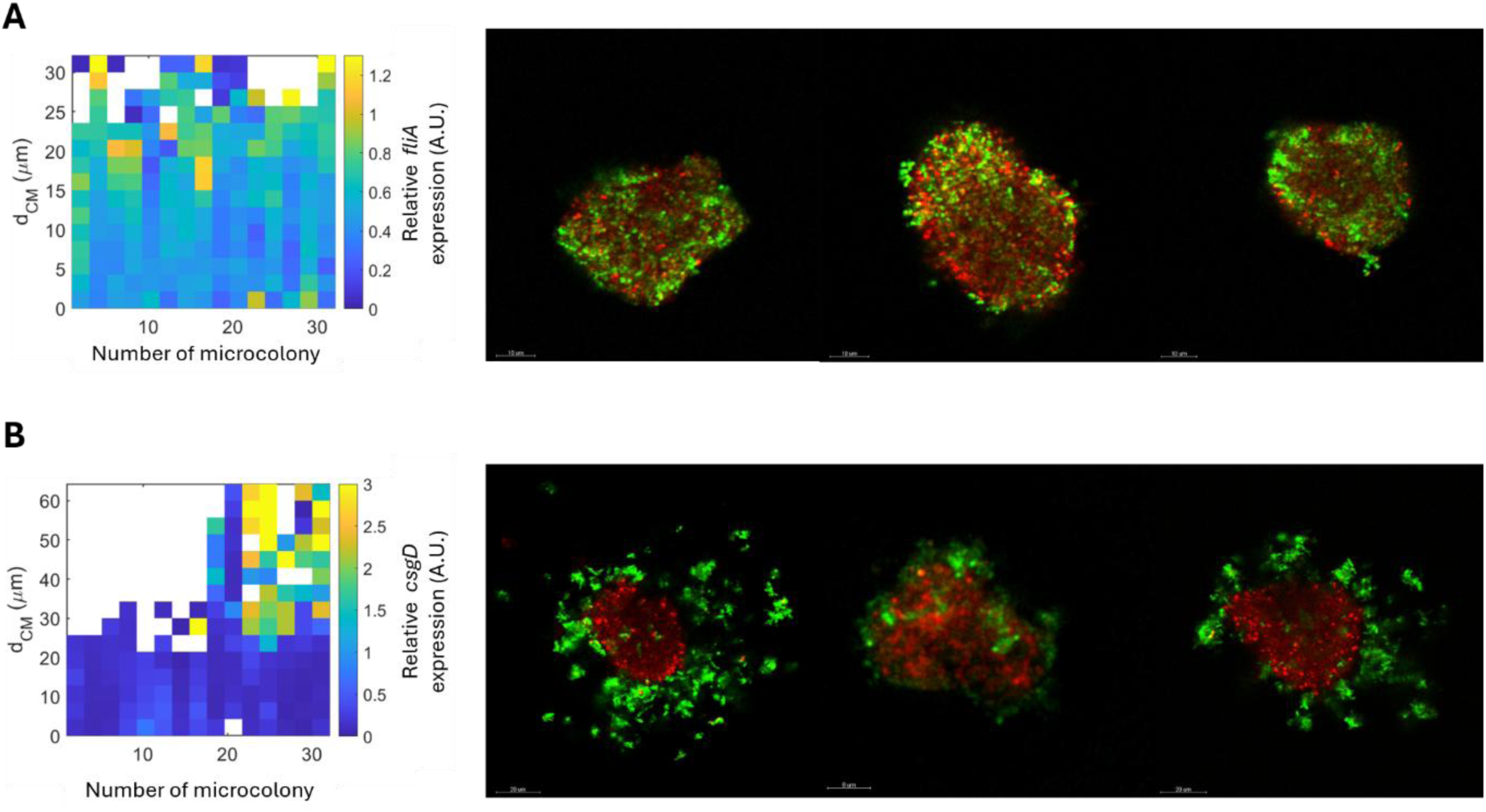
Confocal laser scanning microscopy (CSLM) visualisation of GFP reporter expression in *Salmonella* Typhimurium microcolonies. Four-day-old microcolonies grown at 25 °C in NaCl-free hydrogel (ST) were analysed using green fluorescent protein (GFP) transcriptional reporters. GPP fluorescence (in green) reports promoter activity of **A**: *ffiA* gene, and **B:** *csgD* gene, whereas total bacterial biomass was counterstained with Syto61 (in red). Kymographs show the ratio of green fluorescence intensity over red fluorescence intensity (GFP/Syto61) as a function of the distance from the centre of the microcolony (d_cm_) and are converted into arbitrary units (A.U.) (n = 30). The white spots in the graphs indicate intensity values below the threshold, representing no significant signal for that specific tile. Representative CSLM images, processed in Imaris software, of independent microcolonies are presented for each fluorescent reporter gene (all 30 CSLM images are provided in **Supplementary Figure S13**).

Reporter analyses revealed distinct spatial patterns of promoter activity within NaCl-free microcolonies: *ffiA* promoter activity was broadly distributed but upregulated toward the periphery, whereas *csgD* gene activity was confined to discrete cellular clusters. From the centre to the periphery of the microcolonies, *ffiA* expression was detected throughout the bacterial population, with the highest fluorescent intensity at the periphery, reaching a relative fluorescence of 1.2 (**Figure 7A**). The peripheral enrichment observed with the *ffiA* reporter was consistent with the mean RT-qPCR trend (SFTc versus SFTp, log2FC = −0.72), although substantial variability among RT-qPCR replicates suggested that the magnitude and consistency of this spatial gradient varied among microcolonies. Another flagellar hook-associated gene, *ffgB,* was investigated using fluorescent reporter expression (**Supplementary Figure S14C**). Overall, *ffgB* expression was lower within microcolonies than that observed for *ffiA* and displayed greater variability between microcolonies. Nevertheless, when expression was detected, the fluorescence pattern was generally diffuse throughout the microcolony, with no evident differentiation between the centre and peripheral regions. Accordingly, no reproducible centre-to-periphery gradient in *ffgB* promoter activity, when detectable, could be identified under the conditions tested. Moreover, one biological replicate corresponds to the pool of regions from 15 to 20 microcolonies. In contrast, *csgD* promoter activity was restricted to discrete cellular clusters, preferentially located toward the microcolony periphery, revealing a pronounced spatial heterogeneity in the activation of this biofilm-associated regulatory pathway (**Figure 7B**). Another exopolysaccharide-related gene, *wcaA*, involved in colonic acid biosynthesis, was analysed using a fluorescent reporter (**Supplementary Figure S14D**). In contrast to *csgD* gene activity, no *wcaA* reporter signal above the detection threshold was observed in microcolonies not exposed to 6% NaCl, consistent with the low transcriptional activity inferred from the corresponding RNA-seq condition. However, the transcriptomic analysis indicated upregulation in microcolonies exposed to 6% NaCl. Under osmotic-stress conditions, variable GFP and Syto61 signal quality prevented robust quantitative analysis across the complete dataset. Reporter patterns obtained under these conditions should therefore be considered exploratory and require independent validation. Consequently, these observations require further validation. Finally, as for *ffiA*, RT-qPCR results for the *csgD* gene in the SFTc vs SFTp comparison are highly heterogeneous: log2FC values of 0.58, -2.30, and -0.55, but the mean tends toward downregulation in the centred cells. Nevertheless, for the two genes, the log2FC mean trends toward an upregulation of the genes at the microcolony peripheries. For the other genes studied by LCM and RT-qPCR, transcriptional GFP reporters are not available.

Despite inter-microcolony variability, the reporter data revealed marked microscale heterogeneity in promoter activity. The spatial patterns observed for *ffiA* and *csgD* were broadly consistent with the trends obtained by LCM-RT-qPCR, providing an independent line of evidence that transcriptional states were not uniformly distributed within the microcolonies. In addition to the genes investigated by LCM and RT-qPCR, and those involved in metabolic pathways associated with the sessile lifestyle, a fluorescent reporter targeting the putative gene *STM1851* was also examined. As shown in **Supplementary Figure S14E**, *STM1851* exhibited a highly localised spatial expression pattern within the microcolonies. More specifically, reporter activity was predominantly detected in peripheral cells. The preferential peripheral activity of the *STM1851* reporter suggested that this putative gene could serve as another marker of spatially differentiated cells within the microcolony. Whether this pattern was associated with differences in nutrient availability, oxygenation, or other local environmental parameters remains to be determined. These observations were consistent with physiological specialisation within the bacterial community, with distinct transcriptional programs associated with different spatial positions and local microenvironments.

### 7. Spatial structuring generates physiological signatures consistent with pre-adaptation to digestive constraints

Recent work has demonstrated that growth within food-like structured matrices (hydrogel) generates emergent physiological properties in major foodborne pathogens, including *S.* Typhimurium. These include enhanced growth under osmotic stress and increased survival following simulated gastrointestinal transit (**Hoch et al. 2026**). In contrast to our previous study, which included NaCl during growth, gastrointestinal stresses were applied after growth. The present data, therefore, allowed us to investigate whether the physiological state established during growth in a structured matrix, with or without osmotic stress, could contribute to tolerance to a later digestive challenge. To address this question, we re-examined the transcriptomic, metabolomic, lipidomic, and spatial transcriptomic datasets to identify functions potentially associated with adaptation to digestive stress. Particular attention was given to pathways involved in acid resistance, oxidative stress protection, membrane remodelling and adaptive functions only present within microcolonies.

Several transcriptional and physiological signatures previously identified in spatialised populations were consistent with enhanced resistance to the hostile conditions encountered during gastrointestinal transit. Unlike classical acid resistance systems, which were not enriched in spatialised populations (*speF, potE, adi* or *cadB*), growth within microcolonies was associated with the activation of a broad stress-adaptation programme. The strong induction of *rpoS* (log2FC ST vs PT = 4.8), *dps* (log2FC ST versus PT = 4.2), *bolA* (log2FC ST versus PT = 4.95), and several envelope and protein quality-control functions suggested that spatially structured populations acquire a physiological state characteristic of stress-tolerant and stationary-phase cells. Despite the strong *rpoS* upregulation detected by RNA-seq, no *rpoS* reporter signal above the fluorescence threshold was detected. This discrepancy may reflect differences between the reporter and RNA-seq experimental systems and does not permit conclusions about the spatial distribution of *rpoS* expression from the reporter dataset alone (**Supplementary Figure S14A**). Such a transcriptional state could increase bacterial robustness during subsequent exposure to gastrointestinal constraints encountered during digestion, such as acidity and oxidative stress.

Spatialised populations also displayed enhanced expression of functions associated with oxidative stress resistance. The manganese and iron transport system (*sitABCD*) was strongly upregulated (log2FC ST versus PT ≈ 7) together with pathways involved in inorganic ion transport and metabolism. Because Mn^2+^ and Fe^2+^ homeostasis contributed to protection against reactive oxygen species, these changes were consistent with an increased capacity to withstand oxidative damage. In addition, spatial transcriptomic analyses revealed a tendency toward the upregulation of *katE* gene in peripheral regions of salt-exposed microcolonies, suggesting localised activation of antioxidant defences.

Given the association observed between spatial growth and stress digestive tolerance, changes in the expression of virulence-associated genes, particularly those located in *Salmonella* pathogenicity island SPIs, we further investigated. Among SPI-1 genes, the master regulators *hilA, hilD* and *hilC* were significantly downregulated in microcolonies compared to planktonic cells. Analysis of the *hilA* reporter revealed substantial variability among microcolonies (**Supplementary Figure S14B**). When detectable, *hilA* reporter activity was broadly distributed across the microcolony rather than restricted to a reproducible spatial subpopulation. Likewise, several genes encoding components of the type III secretion system and invasion effectors (*invA, sipABC* and *sopAB*) exhibited reduced expression levels. Together, these results indicated a global repression of the invasion programme in microcolonies, concomitant with the activation of the c-di-GMP second messenger. In contrast, numerous SPI-2 genes were upregulated under spatially structured growth. The regulatory genes *ssrA* and *ssrB,* together with several structural components of the SPI-2 secretion system (*ssa* operon), were significantly upregulated, further supporting a persistence-related transcriptional programme. This virulence-associated transcriptional profile coincided with the induction of stress-response and biofilm-associated functions, suggesting that spatial growth shifts *Salmonella* toward a physiological state displaying reduced invasion-associated transcription and enhanced stress-adaptive features. The concomitant repression of SPI-1 and the induction of SPI-2 genes indicated a marked reorganisation of virulence-associated transcriptional programmes during spatial growth, consistent with a shift away from an acute invasion-associated state.

The spatial heterogeneity observed within microcolonies may also contribute to the enhanced gastrointestinal tolerance. Rather than being homogeneous, microcolonies contain transcriptionally differentiated subpopulations that occupy distinct local microenvironments.

Under osmotic stress, functions involved in stress adaptation, antioxidant defences, and biofilm formation had increased activity in peripheral regions. The presence of these transcriptionally distinct subpopulations may contribute to population-level robustness during subsequent environmental challenges. Moreover, several genes associated with acid stress resistance were upregulated in microcolonies grown in 6% NaCl compared to NaCl-free microcolonies. These included genes in the *cadABC* operon, which encodes lysine decarboxylase to maintain cytoplasmic pH; activation of *the adi* system, the main component of acid pH resistance in *Enterobacteriaceae;* and activation of polyamine metabolism (*speF* and *potE*). Interestingly, exposure of microcolonies to osmotic stress modified the expression of genes involved in SPI, with repression of the SPI-2 regulator and effector, compared to the spatialised NaCl-free condition. This result indicated that NaCl exposure redirects the bacterial response towards osmoadaptation, acid stress preparation, and proton homeostasis.

Finally, rather than inducing a specific response to gastrointestinal stress, spatial growth as microcolonies appeared to generate a diversified community in which some cells already displayed physiological traits associated with resistance to acid, oxidative and membrane-targeting stresses encountered within the gastrointestinal tract. Thus, even in the absence of direct exposure to gastrointestinal conditions, spatial growth established a heterogeneous physiological state containing molecular signatures consistent with enhanced tolerance to subsequent gastrointestinal stresses.

## Discussion

Traditionally, studies investigating the impact of stress on foodborne pathogens have used homogeneous liquid culture models. However, bacteria present in food matrices typically grow in spatially organised communities (Brocklehurst, Mitchell, Smith 1997; Nath et al. 2022; Saint Martin, Darsonval, et al. 2022). Surface biofilms and embedded colonies are increasingly viewed as multicellular bacterial systems possessing emergent properties that cannot be inferred from the behaviour of individual planktonic cells (**Flemming et al. 2016; Hoch et al. 2026**). Our findings identify molecular signatures and candidate mechanisms associated with these adaptive responses through the integration of transcriptomic, metabolomic and lipidomic analyses.

Our results show that spatial growth strongly affects *Salmonella* physiology. In the multi-omics analyses, the main separation was observed between planktonic and spatialised populations, while the effect of NaCl was smaller. DIABLO integration identified coordinated multivariate signatures across the transcriptomic, metabolomic and lipidomic datasets that discriminated the experimental conditions. These observations illustrate the value of integrating complementary omics approaches to obtain a more comprehensive view of bacterial adaptation to environmental constraints (**Rohart et al. 2017**). Indeed, RNA-seq provides information about regulatory networks and metabolic pathways, while ^1^H NMR captures the resulting metabolite phenotype by detecting metabolite depletion or accumulation, and GC-MS characterises changes in membrane lipid composition, particularly fatty acid composition. Overall, spatial organisation exerted a strong influence on bacterial physiology, leading to modifications in gene expression patterns. This transcriptional reprogramming was marked by physiological heterogeneity within three-dimensional communities, highlighting distinct cellular states among bacteria cells within the same microcolony. Similar physiological heterogeneity has been described in biofilms, where gradients of nutrients, oxygen and metabolites generate different cellular states (**Flemming et al. 2016; Jo, Price-Whelan, Dietrich 2022; Stewart, Franklin 2008**). However, our system differs from classical surface biofilms because the bacteria are immobilised within a hydrogel matrix. Nevertheless, local environmental conditions have also been described in bacterial colonies (**Jeanson et al. 2015**). This suggests that physical confinement itself can contribute to the establishment of specific physiological states, as has also been proposed in other structured bacterial systems (**Dufrêne, Persat 2020; Dupont et al. 2026; Monnappa et al. 2026**).

Several transcriptional changes observed in microcolonies were related to functions commonly associated with sessile or biofilm lifestyles. One of the most prominent changes was the upregulation of the master biofilm regulator *csgD,* along with several curli-associated genes (*csgA, csgE,* and *csgG*) and the diguanylate cyclase gene *yaiC*. These genes are known to be central in the *Salmonella* surface biofilm regulatory network and are strongly controlled by c-di-GMP signalling (**Gerstel, Römling 2003; Lamprokostopoulou, Römling 2021; Simm et al. 2014**). The activation of this system was also supported by the downregulation of numerous SPI-1 genes, including *hilA, hilD, sipA* and *sipB*, which are required for epithelial invasion and acute virulence (**dos Santos, Ferrari, Conte-Junior 2019**). The upregulation of fimbriale genes, including *stdA*, suggested that adhesion-associated functions could potentially contribute to the phenotype of spatialised populations (**Shippy et al. 2013**); however, their contribution to the enhanced gastrointestinal survival observed previously remains to be functionally established (**Hoch et al. 2026**). In addition, the activation of the *rpoS, dps,* and *bolA* genes, which are associated with stationary-phase adaptation, stress tolerance, and persistence, was consistent with a sessile lifestyle (**Silva, Galego, Arraiano 2023; Zhao et al. 2026**). Together, these transcriptional changes indicate that growth as matrix-embedded microcolonies promotes a physiological state that shares several features with canonical *Salmonella* biofilm-associated programmes.

Other pathways also constitute a stronger metabolic signature associated with spatial growth. ^13^C NMR has previously been used to correlate the glyoxylate cycle with fatty acid compositions and cell constituents (**Walsh, Koshland 1984**). Nevertheless, ^1^H NMR provides more detailed information on *Salmonella* metabolite production. For example, the activation of the glyoxylate cycle, a metabolic bypass that diverts carbon flux away from the decarboxylation steps of the TCA cycle (**Ahn et al. 2016; Kornberg 1966; Wilson, Maloy 1987**), is also linked to persistence and stress adaptation in *Salmonella*. Mutants lacking *aceA* exhibit reduced long-term survival and impaired persistence in animal models (**Fang et al. 2005**), whereas malate synthase (*aceB*) contributes to resistance against oxidative and nutrient stresses (**Sarkhel et al. 2022**). Extracellular metabolites detected by ^1^H NMR were also associated with transcriptional changes. Planktonic populations accumulated several fermentation products, including acetate, lactate, ethanol, succinate and formate. In microcolonies, some of these metabolites were first produced but later decreased or disappeared from the extracellular medium. Control experiments showed that this disappearance was not due to the hydrogel or to spontaneous degradation, suggesting that the bacteria reutilised these compounds during growth. These results suggest an important reorganisation of carbon utilisation in spatialised populations. **Jeckel et al. 2023**, demonstrated the existence of a spatiotemporal cross-feeding mechanism in *Bacillus subtilis*, in which metabolites (pyruvate and fumarate) secreted by one subpopulation are consumed by cells from subsequent generations. The transient accumulation and subsequent disappearance of these extracellular metabolites are consistent with their reutilisation by spatially localised populations and raise the possibility of spatiotemporal metabolic cross-feeding within *Salmonella* microcolonies, although this mechanism remains to be directly demonstrated.

Another important difference between *Salmonella* lifestyles concerned membrane composition. The lipidomic data revealed an unexpected decrease in WAMT in spatialised populations, which indicated increased membrane fluidity. This observation contrasts with several studies describing biofilm-associated increases in saturated fatty acids and membrane rigidity (**Dubois-Brissonnet, Trotier, Briandet 2016**). Nevertheless, increases in SFA in biofilm appeared to be less present in Gram-negative bacteria. Moreover, another explanation is that fatty acid measurements represent averages across thousands of microcolonies, which were pooled to achieve sufficient biomass. Given the observed heterogeneity within individual microcolonies, these pooled measurements may have limited differences between coexisting subpopulations. The response to NaCl was also different between the two lifestyles. In planktonic populations, NaCl strongly increased the proportion of saturated fatty acids, whereas the lipid profile of spatialised populations changed less and remained closer to the spatialised control condition. Changes in membrane fatty acid composition, and more specifically, rigidification of the membrane (an increase in saturated fatty acids) to prevent plasmolysis and maintain intracellular homeostasis, are known to contribute to bacterial adaptation to environmental stresses (**Sleator, Hill 2002**). In *Salmonella*, modifications in membrane fatty acid composition in response to previous growth conditions have also been associated with changes in stress resistance (**Álvarez-Ordóñez et al. 2008**). However, additional experiments would be required to determine whether these membrane changes directly contribute to the different responses to osmotic stress.

An important result of this study is that spatial growth altered the physiological response of *Salmonella* during growth under osmotic stress. Planktonic populations grown in the presence of 6% NaCl exhibited substantial transcriptional and metabolic differences compared with planktonic populations grown under control conditions. In contrast, the differences between spatialised populations grown with or without NaCl were more limited. Both lifestyles activated classical mechanisms of osmotic adaptation, but spatial growth established a distinct physiological background that modified the response developing in the presence of NaCl. We therefore propose that spatial organisation not only increases osmotic protection but also alters how *Salmonella* adapts as it grows under osmotic stress. The spatial analyses also showed that a microcolony is not a homogeneous population: RT-qPCR following LCM revealed differences between central and peripheral regions, although substantial variability was observed among biological replicates. Fluorescent reporters further revealed gene-specific spatial patterns within microcolonies, with broadly distributed but peripherally enriched *ffiA* promoter activity and *csgD* activity restricted to discrete cellular clusters. Similar heterogeneity has been described in bacterial biofilms and macrocolonies, where gradients of nutrients, oxygen and metabolic products generate different physiological states (**Jo, Price-Whelan, Dietrich 2022; Serra, Hengge 2014; Stewart, Franklin 2008**). Overall, these spatial expression patterns are consistent with physiological differentiation along the centre-to-periphery axis of the microcolonies. Peripheral regions preferentially expressed genes associated with adhesion, motility and membrane adaptation, including *csgD, ffiA* and *fabA,* whereas central regions tended to show higher expression of selected stress-associated functions. Although such patterns are compatible with spatial differences in nutrient availability, oxygenation and metabolite accumulation, these local physicochemical parameters were not directly measured in the present study.

Unexpectedly, *ffiA* and *csgD* displayed broadly similar spatial expression patterns. As these genes are generally associated with antagonistic lifestyles, motility and sessility, so their co-occurrence in peripheral cells appears counterintuitive (**Lamprokostopoulou, Römling 2021**). However, this observation may be explained by the use of relatively long-lived GFP (**Robijns et al. 2014**). Thus, the fluorescence observed after four days of growth likely integrates successive transcriptional states experienced by the cells, during microcolony development, rather than providing a snapshot of current gene expression as LCM and RT-qPCR experiments. Temporal transitions between motile and sessile lifestyles may therefore be obscured by the persistence of the GFP signal.

In *Salmonella*, bistable expression of *csgD* has also been shown to generate distinct subpopulations associated with different physiological functions (**Grantcharova et al. 2010; MacKenzie et al. 2015**). Spatial transcriptional heterogeneity has also been directly demonstrated across different *Bacillus subtilis* biofilm communities (**Dergham et al. 2023**). Our results suggest that this type of heterogeneity can also develop when the spatial organisation of the population is imposed by the surrounding matrix. Because only a limited number of genes were analysed by LCM and RT-qPCR, this approach should be considered as a targeted spatial gene-expression analysis rather than a complete spatial transcriptomic analysis. Higher-resolution approaches will be useful to better characterise these different subpopulations. Other techniques have been developed in recent years to decode the spatiotemporal heterogeneity of complex microbial communities (**Alves, Pombinho, Cabanes 2026; Dar et al. 2021; Espinoza Miranda et al. 2025**).

Multi-omics approaches are highly valuable for elucidating the complexity of bacterial adaptation in spatially organised *Salmonella* communities growing in hydrogels. However, each approach remains subject to technical and biological limitations: gene expression, metabolite production and fatty acid composition were measured at a single time point during bacterial growth; lower quality in ^1^H NMR spectra was attributed to the low concentration of metabolites in the samples and the use of water as solvent; fatty acid profiles were obtained from pooled bacterial populations, which prevented individual variability from being taken into account; the genetic material recovered by LCM was limited in quantity and variable in quality. To date, LCM has been primarily applied to eukaryotic tissues and cells, with only limited use in prokaryotic systems (**Lenz et al. 2008; Pérez-Osorio, Williamson, Franklin 2010; Podgorny, Lazarev 2017**). The present study, therefore, constitutes a proof of concept, demonstrating the strong potential of this technique for investigating the spatial organisation of bacterial communities. Finally, despite these limitations, the different techniques offer opportunities for methodological improvements that should be considered when interpreting the results and designing future studies.

Finally, these results can help to understand the gastrointestinal cross-protection observed in our previous study (**Hoch et al. 2026**). Spatialised populations showed upregulation of several genes associated with general stress responses, including *rpoS*, *dps* and *bolA*, along with changes in membrane composition and oxidative stress-related functions. Under NaCl stress, additional changes were observed in pathways involved in acid adaptation and proton homeostasis. This transcriptional reprogramming provides a plausible mechanistic basis for the cross-protection observed between osmotic adaptation and digestive tolerance. Moreover, catalase activity associated with *katE* has previously been described as protective against oxidative stress (**Benoit, Maier 2016**). These changes are compatible with a physiological state that could favour survival during subsequent gastrointestinal stresses. However, the present data do not demonstrate that these pathways are directly responsible for the previously observed increase in gastrointestinal tolerance. Functional validation of selected pathways will therefore be necessary. Interestingly, heterogeneous *csgD* expression in *S. Typhimurium* has previously been proposed to connect bacterial states associated with virulence and persistence (**MacKenzie et al. 2015**), while c-di-GMP signalling has been implicated in the balance between biofilm formation and virulence-associated properties (**Ahmad et al. 2011**).

Overall, our results suggest that growth in a spatially structured environment establishes a physiological state that is clearly different from the planktonic state. This involves changes in carbon metabolism, membrane composition, stress-response functions and the appearance of transcriptional heterogeneity within microcolonies. Importantly, this physiological state also modifies how *Salmonella* responds to osmotic stress during growth. Together, these findings support the view that spatial organisation is not merely a structural feature of bacterial growth, but an environmental determinant that shapes physiological diversification and the subsequent capacity of foodborne pathogens to adapt to stress.

## Materials s Methods

### Bacterial strains and spatial growth conditions

*Salmonella enterica* subsp. Enterica serovar Typhimurium LT2 strain was used throughout this study and grown in tryptone-soya broth (30 g/L) supplemented with yeast extract (6 g/L) (ThermoFisher, Dardilly, France) at 25 °C. Growing medium for planktonic and hydrogel cultures was prepared as previously described in **Hoch et al. 2026**. *Salmonella enterica* serovar Typhimurium ATCC 14028 wild strain was used for reporter fusion constructions. Fluorescent fusion reporters were constructed in **Robijns et al. 2014**. All bacteria containing a reporter fusion plasmid were inoculated in TSBYe supplemented with 100 µg/mL of ampicillin and incubated overnight at 37 °C.

All planktonic assays were performed at 25 °C in 15 mL Eppendorf tubes in TSBYe, with or without 6% NaCl supplementation. For spatialised assays, a hydrogel composed of half low-melting-point agarose (final concentration: 0.5% agarose) and half of either a doubled-concentration TSBYe control or a 6% NaCl medium was grown in 96-well plates (200 µL) and incubated at 25 °C. For transcriptomic analysis (RNA-seq and RT-qPCR), media were supplemented with 1 mg/L calcofluor white (Merck, Darmstadt, Germany).

The samples shall be named as follows: PT for planktonic cultures without NaCl, PNaCl for planktonic cultures supplemented with 6% NaCl, ST for spatialised cultures without NaCl and SNaCl for spatialised cultures supplemented with 6% NaCl.

### Sample collection for RNA-seq and GC-MS analyses

For liquid samples: for each condition, from a 4-day culture, 4 mL is centrifuged at 12,000 x g at 4 °C for 2 minutes. The pellet is washed with 1 mL of TE (10 mM Tris, 1 mM EDTA, pH 8) + 60 µL of 1 M EDTA. A centrifugation step for 2 minutes (12,000 x g at 4 °C) to discard the supernatant was performed, and samples were snap-frozen in liquid nitrogen for transfer to -80 °C for RNA and GC-MS extraction.

For spatialised samples: 20 wells are pooled per condition into a 50 mL Falcon tube, yielding a final hydrogel volume of 4 mL. Approximately 20 glass beads (2 mm) were added, along with 20 mL TE (10 mM Tris, 1 mM EDTA, pH=8) + 1,2 mL 1 M EDTA, and vortexed at maximum speed for 10-20 seconds. Five millilitres of this mix were added to a 20 mL syringe equipped with two Whatman paper discs (category 4, pore size 20-25 µm). After filtration, the filters were removed and replaced with new ones. The filtration step was repeated until the entire mix had been filtered. The Falcon tube was rinsed with 2 mL TE + 120 µL EDTA and filtered as well. The filtrate was aliquoted into 2 mL Eppendorf tubes and centrifuged for 1 minute (12,000 x g at 4 °C) to discard the supernatant. Samples were snap-frozen in liquid nitrogen and transferred to -80 °C for RNA and GC-MS extraction.

### Metabolome extraction for NMR analysis

For liquid cultures: for each condition, 3 mL of 4-day-old cultures were centrifuged at 5,000 x g for 5 min at 4 °C. The resulting supernatants were filtered through 0.22 µm PES membranes and stored at 4 °C until NMR analysis, which was carried out on the day of sample collection. Three biological replicates were done per condition, each one in three technical replicates.

For spatialised cultures: 15 wells are pooled into a 15 mL Eppendorf tube, yielding a final hydrogel volume of 3 mL per condition. Approximately 10 glass beads (2 mm) were added, along with 3 mL of physiological water (9 g/L NaCl), and the mixture was vortexed at maximum speed for 10-20 seconds. Samples were centrifuged at 5,000 x g for 5 min at 4 °C. The resulting supernatants were filtered through 0.22 µm PES membranes and stored at 4 °C until NMR analysis, which was carried out on the day of sample collection. Three biological replicates were done per condition, each one in three technical replicates.

To monitor metabolite dynamics over the 4-day growth period, planktonic cultures and hydrogels were sampled at successive growth stages (Days 1, 2, 3 and 4). Metabolites were extracted using the same protocol as for endpoint analyses and stored at 4 °C until analysis on the same day.

Deuterated water (D_2_O, D ≥ 99.9 %) containing 0.05 % (w/w) 3-(trimethylsilyl)propionic acid-2.2.3.3-d_4_ sodium salt (TSP) was obtained from Merck-Sigma Group (St. Louis, MO, USA). For planktonic cultures, 500 µL of the prepared sample and 500 µL of D_2_O containing the reference standard TSP were mixed using a Vortex Genie 2 (Bender C Hobein AG, Zurich, Switzerland) at room temperature. For spatialised cultures, the sample-to-D_2_O ratio is 3:1. Finally, 600 μL of the mixture was transferred into a 5mm NMR tube for ^1^H NMR analysis.

### RNA extraction for RNA-seq

Pellets were recovered from -80 °C and suspended in 1 mL TRIzol reagent. The RNA extraction and cleanup protocols, as described by, **Dergham et al. 2023** were applied. Nanodrop and Bioanalyzer instruments were used for quantity and quality controls. Library preparation, including ribosomal RNA depletion and sequencing, was performed by the I2BC platform (Gif-sur-Yvette, France) using TruSeq Total RNA Stranded and Ribo-Zero Bacteria Illumina kits, and an Illumina NextSeq 2000 PE to generate paired-end reads (2 x 50 bp).

### RNA-seq data analysis

Primary data processing was performed by the I2BC platform and consisted of demultiplexing (with bcl-convert 4.4.6), adapter trimming (Cutadapt 4.7), quality control (FastQC v0.11.5), and mapping (with bwa 0.7.18) against the *Salmonella enterica* Typhimurium LT2 genome (NCBI GenBank: AE006468.2). This generated between 23 M and 54 M uniquely mapped reads per sample, which were summarised as read counts for 4439 genes (featureCounts 2.0.8).

The downstream analysis was performed using R version 4.3.2. DEGs (Differentially Expressed Genes) were detected using the R package *DESeq2* (version 1.46.0) to estimate p-values and log2-foldchanges (log2FC). PCA (Principal component analysis) and a volcano plot were obtained using the *DESeq2* package. Genes identified by RNA-seq were functionally annotated by assigning them to COG (Clusters of Orthologous Groups) categories on the basis of protein homology. Subsequently, each gene was mapped to its corresponding KEGG pathway entry, allowing integration into the metabolic and functional pathways curated in the KEGG database. All graphics (heatmaps or bubble charts) were constructed using *Tidyverse, clusterProfiler* and *ggplot2* R packages (version 4.3.2). The whole transcriptomic dataset has been deposited in GEO (accession number <u>GSE342574</u>).

### Membrane fatty acids analysis

Extraction and methylation of fatty acids (FA) were carried out directly on bacterial pellets as previously described (Dubois-Brissonnet, Trotier, Briandet 2016; Touche et al. 2023). Whole-cell FA were saponified and esterified by first methanolic NaOH (addition of 1 mL of 3.75 M NaOH in 50% (v/v) methanol to the pellet and incubation at 100 °C for 30 min) and then by methanolic HCl (addition of 2 mL of 3.25 M HCl in 45 % (v/v) methanol solution and incubation at 80 °C for 10 min). FA methyl esters (FAME) were extracted with a 1:1 (v/v) diethyl ether/cyclohexane solution. At the end, the organic phase was washed with dilute base (0.3 M NaOH).

Analytical gas chromatography of FAME was carried out on a GCMS Trace 1300/ISQ 7000 system (Thermo Fisher Scientific) equipped with a BPX70 capillary column (25 m, 0.22 mm internal diameter; SGE, Victoria, Australia). The column temperature was set to 100 °C for 1 min, then increased to 170 °C at 2 °C/min. FA species were identified using MS databases (Replib, Mainlib, FAME2011). Fatty acids were grouped in classes: saturated fatty acids (SFA), unsaturated fatty acids (UFA), hydroxylated fatty acids (HFA) and cyclopropane fatty acids (CFA). Results are the average of at least 9 profiles (three injections of three extractions from independent cultures) per condition. The relative abundance of FA species was expressed as the percentage of the total FAME peak area. An additional indicator of membrane adaptation was also calculated, namely, weighted-average melting temperature (WAMT). WAMT is calculated from the melting temperature of each individual FA and its ratio in the membrane composition.

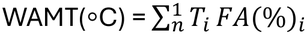

Where T_i_ (°C) is the melting temperature of FA_i_, FA (%) is the percentage of FA_i_, and FA_1_ to FA_n_ are each of the FAs present in the membrane composition. Melting temperatures of FA (°C) were obtained from CAS Common Chemistry (<u>CAS Common Chemistry</u>).

### Metabolome analysis by NMR

^1^H NMR spectra were obtained using a Bruker Avance III HD at 9.4 Tesla (Bruker Biospin, Rheinstetten, Germany) equipped with a PA BBO 400S1 BBF-H-D-05 Z SP probe operating at a proton Larmor frequency of 400 MHz. The acquisition was performed using the TOPSPIN online software (version 3.5 pl6, Bruker Biospin, Rheinstetten, Germany). All spectra were acquired without rotation (spin-off conditions) to avoid undesired spinning bands and at a temperature of 293.0 ± 0.1 K. ^1^H NMR spectra were obtained using the Bruker one-dimensional NOESY gradient water suppression pulse sequence (noesygppr1d). Concerning the acquisition conditions, the values of the number of scans (NS) and the receiver gain (RG) were respectively 184 and 128 and the other conditions were: spectral width (SW) = 21 ppm, 8417 Hz; transmitter frequency (O1P) = 4,7 ppm, 1881 Hz; size of FID (TD) = 67338; relaxation delay (D1) = 1 s; acquisition time (AQ) = 4 s; FID resolution = 0.25 Hz.

The duration of the ^1^H NMR experiment was about 16 min per sample. All Free Induction Decay (FID) signals were then multiplied by an exponential function with a line broadening factor of 0.3 Hz prior to Fourier transformation. Once ^1^H NMR spectra were obtained, they were baseline-corrected, phased, and calibrated to the TSP signal using TOPSPIN offline software (version 4.5.0). Chemical shifts (δ) were expressed in parts per million (ppm) and relative to the TSP signal (δ 0.0 ppm). The whole ^1^H NMR dataset has been deposited in nmrXiv (https://doi.org/10.57992/nmrxiv.e3b5c3e7).

The ^1^H NMR pre-processed data were treated using the online website NMRProcFlow v1.4. Spectra were binned from 0.1 to 4.80 ppm and from 4.90 to 10.00 ppm using a 0.04 ppm bin width. A signal-to-noise ratio (SNR) threshold of 3 was used, meaning that all bins with SNR < 3 were removed. The 4.80-4.90 ppm region was excluded in order to remove residual signals arising from water presaturation. A total of 214 bins were obtained, and the resulting matrix was exported as a CSV file for subsequent statistical analyses.

### Laser capture microdissection

After 4 days of growth, either with or without 6% NaCl supplementation, a hydrogel sample was collected and fixed overnight at 4 °C in a 75:25 EAA solution (absolute ethanol and glacial acetic acid). After fixation, 3 washing steps were performed in RNase-free water at 4 °C for 1 hour with shaking. The hydrogels were embedded in Tissue-Tek O.C.T. compound and snap-frozen in liquid nitrogen, then stored at -80 °C for cryosectioning. Microcolony sections (20 µm thick) were obtained by sectioning the solidified O.C.T.-containing microcolonies in a cryostat (FSE Shandon, ThermoFisher, France) at -20 °C and in an RNase-free environment (material treated with RNase-free reagents and 70% ethanol). For each biological replicate and to obtain enough material for downstream analyses, 10 cross-sections per hydrogel were gathered on Arcturus PEN membrane frame RNase-free slides (Excilone, France). Sections were fixed immediately in 75% cold ethanol and coloured by the following protocol: a homemade 1% cresyl violet solution for 20-30 seconds, then 75% ethanol for 30 seconds to rinse off excess cresyl violet and fix the marking. The PEN Membrane Frame Slides were assembled in a “sandwich” configuration according to the setup illustrated in **Supplementary Figure S15**. To locate the targeted structures, sections were analysed under the optical microscope using the XT® Arcturus Technologies microdissection system (Excilone, France).

The LCM process was carried out using the XT® Arcturus Technologies microdissection system and software. Capture was performed using Arcturus CapSure® LCM Macro Caps (Excilone, France). Microcolonies present inside the hydrogel were identified using the 40X objectives. PEN Membrane Frame Slides allowed the use of standard IR and UV lasers. A UV laser was used to cut around and inside the microcolonies, physically separating the periphery from the centre, using calcofluor fluorescence as a guide, since the fluorescence signal is expressed only at the periphery. An IR laser was used to capture and adhere the samples to the macro caps without contaminating the non-target region. To obtain enough material (at least 100 pg/µL RNA) for downstream analysis, ∼10 microcolonies regions were captured by caps, and 2 caps per biological replicate were stored at -80 °C.

Total RNA was extracted using the Picopure^TM^ RNA Isolation kit (Excilone, France). The quality and quantity of the RNA were assessed using an Agilent 2100 Bioanalyzer (Agilent Technologies, Palo Alto, CA) with an RNA 6000 Pico Lab Chip and analysed with the RNA Integrity Number (RIN) algorithm.

### Pre-amplification and RT-qPCR

For qPCR analysis, a panel of 10 targeted genes, including 8 genes of interest and 2 reference genes (**Table 1**) was selected to measure the differences in gene expression levels between core and peripheral regions of colonies. Primer design was performed using Primer Express 3.0 software (Thermo Fisher Scientific) based on gene-specific sequences identified through the previous global transcriptome analysis. Gene sequences were obtained from *Salmonella enterica* Typhimurium LT2 genome on NCBI (GenBank: AE006468.2).

Primer efficiencies were first tested on non-precious RNA samples (1 µg) extracted from whole *S.* Typhimurium LT2 microcolonies. Reverse transcription (RT) was performed on 16 µL of RNA essentially using the SuperScript^TM^ IV VILO^TM^ (SSIV VILO) Master Mix kit according to the manufacturer’s instructions (Invitrogen, Thermo Fisher Scientific), under the following conditions: 25°C for 10 minutes, 50°C for 10 minutes and 85°C for 5 minutes. Total volume of cDNA was diluted 1000-fold prior to qPCR analysis. qPCR runs were performed using SYBR^TM^ Master Mix PCR Power SYBR^TM^ Green (Thermo Fisher Scientific) according to the manufacturer’s instructions on a QuantStudio 12K Flex thermocycler (Thermo Fisher Scientific). After the optimisation of qPCR systems (efficiency ranging from -3.25 to -3.45), amplification reactions were run in triplicate under the following conditions: 95°C for 15 minutes, 40 cycles of 95°C for 15 seconds and 60°C for 1 minute. The mean cycle threshold (Ct) value for each triplicate was calculated for each gene.

To enable the quantification of 10 genes from very low quantities of RNA obtained after microdissection (between 19 pg and 950 pg), a targeted RNA preamplification step was performed using the TaqMan^TM^ PreAmp Master Mix (Thermo Fisher Scientific), according to the manufacturer’s instructions. The kit was selected to enable target-specific preamplification using gene-specific primers without probes, followed by gene expression quantification using SYBR Green-based qPCR. The gene-specific assays corresponding to the 10 target genes were pooled prior to the preamplification reaction, as recommended by the manufacturer. Briefly, 10 µL of each primer (20 in total) at a concentration of 18 µM were combined to generate a single assay preamp pool for a total volume of 200 µL. Then, the pooled primers were diluted using nuclease-free water (Qiagen) so that each assay is at a final concentration of 0.2X. This preamp pool was used for the preamplification reaction.

To validate the preamplification procedure before applying it to microdissected samples, we performed a linearity test on the reference RNA. This RNA was used at dilutions of 1 and 0.1 ng for preamplified cDNA synthesis to validate the linearity of preamplification of each gene, independently of input amount. The protocol for reverse transcription, followed by the target-specific preamplification procedure consisting of 14 cycles using multiplex assays. In parallel, the corresponding RNA sample was reverse transcribed using the SuperScript^TM^ IV VILO^TM^ (SSIV VILO) Master Mix kit according to the manufacturer’s instructions (Invitrogen, Thermo Fisher Scientific), and analysed without preamplification. The preamplified and non-preamplified cDNA samples were then analysed by qPCR for the 10 target genes. For each gene, the difference between the Ct values of the preamplified sample and the corresponding non-preamplified control was calculated (ΔCt). The ΔCt values obtained for the 10 genes were compared to assess whether the preamplification procedure introduced a gene-specific amplification bias. Once the preamplification procedure was validated using diluted reference RNA, the same protocol was applied to the microdissected samples, which contained only a few picograms of RNA and were therefore particularly limited in quantity. The input amount used for preamplification was 200 pg except for SFNaClp R1, SFTp R2 and SFNaClp R3, for which the RNA amount was not detectable by the bioanalyser.

RT-qPCR data analyses were performed using the R packages: t*idyverse, ggplot2,* and *dplyr*. Reference housekeeping genes *rpoE* and *dnaN* were used for accurate normalisation of data. R codes for analyses are accessible *via* INRAE forge: https://forge.inrae.fr/elodie.hoch/code_availability_multi-omics.git.

### GFP reporter and confocal laser scanning microscopy (CLSM)

Microscopic observations were performed with a Leica Mica Widefield Microhub when calcofluor white staining and with a Leica HCS-SP8 confocal laser scanning microscope for GFP reporter observations, at the INRAe MIMA2 imaging platform (https://mima2.jouy.hub.inrae.fr/). The presence of cellulose inside microcolonies was assessed using calcofluor white dye (blue) and SYTO 61 (red) counterstain (1:500 dilution in TSBYe medium from a stock solution at 5 µM in DMSO (Invitrogen, France)), a nucleic acid dye. For GFP-reporter visualisation, the same counterstain was used. On Leica HCS-SP8 CLSM, SYTO 61 excitation was performed at 633 nm using an HeNe laser, and the emitted fluorescence was recorded at 640-690 nm using a photomultiplier detector. GFP excitation was performed at 485 nm, and the emitted fluorescence was recorded at 501 nm. UV fluorescence of calcofluor white was excited at 355 nm, and the emitted fluorescence was recorded at 430 nm. For 3D stack analysis, a 1 µm step was used between z levels. Image analysis was processed using IMARIS 9.3.1 software (Bitplane, Switzerland). Background fluorescence was first subtracted from all pictures before applying a Gaussian filter to each fluorescence. Kymograms reporting the spatial analysis of fluorescent reporter gene expression in microcolonies were performed using BiofilmQ (version 1.0.1). BiofilmQ image segmentation was performed with cubes of 10 vox. The absence of fluorescence gradients in *Salmonella* wild strains was assessed with a Syto61 biomass dye.

### Statistical analysis

For NMR analysis, the generated data were processed using the online website MetaboAnalyst v6.0, where they were normalised by sum and auto scaled prior to principal component analysis (PCA).

For PCA, omics datasets (transcriptomic, metabolomic and lipidomic) were integrated to perform a global principal component analysis (PCA) using the *prcomp* function from the FactoMineR package (R version 4.3.2). Prior to integration, raw RNA-seq counts were normalised using the DESeq2 pipeline; metabolomic data were normalised and autoscaled; and lipidomic data were log-transformed, centred, and scaled. The resulting normalised matrices were then concatenated into a single multi-omics matrix. Relationships among samples were projected onto the first two principal components, and individuals were visualised in the PC1/PC2 factor plane. Contributions to the PC were subsequently extracted to identify the genes, metabolites, and lipids that most strongly contributed to the overall variability among samples. R codes for analyses are accessible *via* INRAE forge: https://forge.inrae.fr/elodie.hoch/code_availability_multi-omics.git.

For multi-omics data integration, the DIABLO multi-omics integration method from the mixOmics v6.8.5 R package was used. Transcriptomic (RNA-seq), metabolomic (^1^H NMR) and lipidomic (GC-MS fatty acid profiles) datasets were integrated to identify coordinated molecular signatures associated with lifestyle and osmotic stress. Prior to integration, each dataset was centred and scaled to unit variance using the *scale()* function in R. A sparse multi-block partial least squares discriminant analysis (PLSDA) was performed. A design matrix of 0.1 was used to focus primarily on discrimination between environments, leading to the selection of 15 genes, 15 metabolites, and 3 fatty acid families. The final DIABLO model was fitted using the optimal *keepX* values selected during tuning. Sample clustering was visualised using the individual score plots, while relationships among omics layers were assessed using latent component correlations. Multi-omics interactions were explored using a clustered image map (CIM) and relevance networks (with a correlation threshold of r = 0.7). R codes for analyses are accessible *via* INRAE forge: https://forge.inrae.fr/elodie.hoch/code_availability_multi-omics.git.

## Data availability

The whole transcriptomic dataset generated in this study has been deposited in GEO (accession number <u>GSE342574</u>). The whole metabolomic dataset generated in this study has been deposited in nmrXiv (https://doi.org/10.57992/nmrxiv.e3b5c3e7). The R codes used for this study are accessible *via* INRAE forge: https://forge.inrae.fr/elodie.hoch/code_availability_multi-omics.git. The source data document accompanying this paper contains the raw data used to generate Figures 1, 2, 3, 4, 5, and 6. The online version contains supplementary datasets.

## Acknowledgements

This research was funded by Aerial, INRAE, the French National Research Agency (ANR) through the France 2030 initiative (ANR-22-GDFF-000) and “Association Nationale de la Recherche et de la Technologie” (contract 2023/0850). We thank the MIMA2 imaging platform (Microscopie et Imagerie des Microorganismes, Animaux et Aliments, https://doi.org/10.15454/1.5572348210007727E12, for the Leica SP8-HCS microscopy observations. We acknowledge the sequencing and bioinformatics expertise of the I2BC High-throughput sequencing facility, supported by France Génomique (funded by the French National Program “Investissement d’Avenir” ANR-10-INBC-09).

Grammarly was used solely for language and grammar editing. The authors take full responsibility for the content of the manuscript.

## Author contributions

E.H., L.F.O., and R.B., conceptualised the overarching aims of the research study. E.H., Y.D., A.P., H.S., D.W., B.H., C.B., L.F.O., and R.B., conceived and designed the experiments. E.H., Y.D., A.P., J.D., F.D.B., J.L., and C.B., performed the experiments and data acquisition. E.H., Y.D., A.P., J.D., F.D.B., J.L., C.B., L.F.O., and R.B., analysed and interpreted the data. L.F.O., and R.B. had both management and coordination responsibilities for the execution of the research work. L.F.O., and R.B. contributed to the acquisition of the financial support and resources leading to this publication. All authors substantially (i) contributed to the conception or design of the work or the acquisition, analysis or interpretation of the data, (ii) revising critically the manuscript for important intellectual content and (iii) approved the final completed version.

